# Genetic variation for plant growth traits in a common wheat population is dominated by known variants and novel QTL

**DOI:** 10.1101/2020.12.16.422696

**Authors:** Noah DeWitt, Mohammed Guedira, Edwin Lauer, J. Paul Murphy, David Marshall, Mohamed Mergoum, Jerry Johnson, James B. Holland, Gina Brown-Guedira

## Abstract

Genetic variation in growth over the course of the season is a major source of grain yield variation in wheat, and for this reason variants controlling heading date and plant height are among the best-characterized in wheat genetics. While the major variants for these traits have been cloned, the importance of these variants in contributing to genetic variation for plant growth over time is not fully understood. Here we develop a biparental population segregating for major variants for both plant height and flowering time to characterize the genetic architecture of the traits and identify additional novel QTL. We find that additive genetic variation for both traits is almost entirely associated with major and moderate-effect QTL, including four novel heading date QTL and four novel plant height QTL. *FT2* and *Vrn-A3* are proposed as candidate genes underlying QTL on chromosomes 3A and 7A, while *Rht8* is mapped to chromosome 2D. These mapped QTL also underlie genetic variation in a longitudinal analysis of plant growth over time. The oligogenic architecture of these traits is further demonstrated by the superior trait prediction accuracy of QTL-based prediction models compared to polygenic genomic selection models. In a population constructed from two modern wheat cultivars adapted to the southeast U.S., almost all additive genetic variation in plant growth traits is associated with known major variants or novel moderate-effect QTL. Major transgressive segregation was observed in this population despite the similar plant height and heading date characters of the parental lines. This segregation is being driven primarily by a small number of mapped QTL, instead of by many small-effect, undetected QTL. As most breeding populations in the southeast U.S. segregate for known QTL for these traits, genetic variation in plant height and heading date in these populations likely emerges from similar combinations of major and moderate effect QTL. We can make more accurate and cost-effective prediction models by targeted genotyping of key SNPs.

## 1 Introduction

Wheat is a major food crop, contributing nearly 20% of human calories and protein (FAO, 2020). Wheat yield is highly polygenic, with variation in yield emerging from variation in other phenotypes each with different genetic bases. Plant growth traits such as heading date (when the spike emerges from the flag leaf) and adult plant height affect yield by both altering resource partitioning between tissues and changing how plants experience environmental factors. A plant’s height on a given date alters the physical position of the plant within its environment, influencing that plant’s interactions with environmental factors like wind, weed competitors, and rain-splashed pathogens. Differences in heading date change the temporal position of plants at a given developmental stage, exposing them to different weather conditions and disease pressures. Wheat breeders typically select for optimal values of plant height and heading date for a given environment and production system in early generations based on unreplicated head rows. Beyond this selection, improvements in yield resulting from modern plant breeding programs have largely been generated without considering its underlying genetic architecture, including the dependence of final plant yield on variation in plant growth trajectories. Understanding the plant development factors that generate genetic variation in yield is critical to increasing the rate of genetic gain in wheat.

Allelic variation affecting core flowering time genes is strongly associated with the geographic distribution of wheat cultivars, and permits the cultivation of wheat in a wide range of environments. Winter wheat is sown in the fall, when it germinates but maintains the shoot apical meristem beneath the ground to prevent freeze damage. After the accumulation of signal through the vernalization (cold hours), photoperiod (night length), and earliness-per-se (plant age) pathways, plants release from winter dormancy and transition to reproductive development. Allelic series in the *Vernalization1* (*Vrn1*) loci on the three chromosome 5 homeologues condition a spring or winter growth habit by controlling the sensitivity of plants to vernalization (Fig. 1) (Yan et al., 2004; Fu et al., 2005; Li et al., 2013). Additional alleles at these loci, some associated with copy number variation, may also modulate vernalization response in vernalization-sensitive winter lines (Fu et al., 2005; Díaz et al., 2012; Guedira et al., 2014, 2016; Kippes et al., 2018). *Photoperiod1* (*Ppd1*) is another core flowering time gene which integrates signals due to length of nights and allows plants to time flowering based on photoperiod. Variants affecting homeologous *Ppd1* loci on all three genomes lead to constitutive over-expression of *Ppd1* and a photoperiod-insensitive, earlier flowering habit (Beales et al., 2007; Nishida et al., 2013; Guedira et al., 2016). Breeding for an optimal heading date for a given environment allows plants enough time to add additional spikelets per spike prior to heading, which increases grain number, and to accumulate carbohydrates and fill grain, which increases grain size. Non-optimal flowering can expose plants to temperatures below freezing early in the season, or to excessive heat and drought late in the growing season. In southern U.S. field sites, *Vrn1* and *Ppd1* alleles have strong effects on final grain yield of winter wheat (Addison et al., 2016).

**Figure 1:**
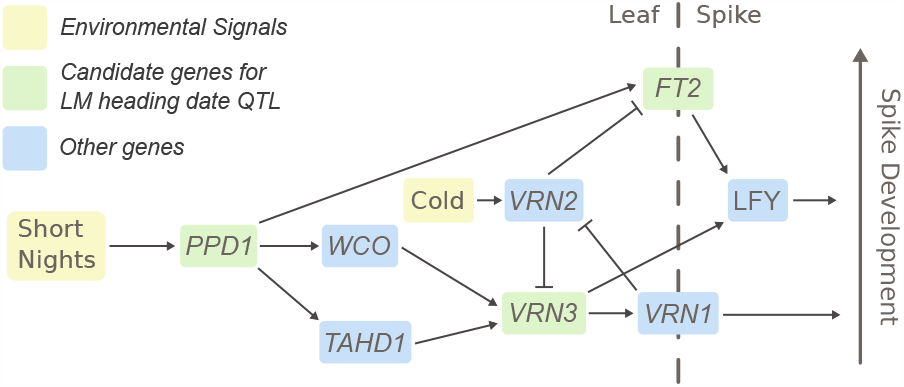
Overview of the wheat flowering time pathway. The gene network through which wheat plants receive and integrate signal about environmental conditions to determine heading date are outlined. Other, intermediate genes are not shown. Genes proposed as candidate for heading QTL in this population are highlighted in green. Other important genes in the flowering time pathway are highlighted in blue; *Wheat CONSTANS (WCO), Triticum aestivum HD1 (TaHD1), VERNALIZATION1 (VRN1), VERNAL-IZATION2 (VRN2)*, and *LEAFY (LFY*).

Introduction of “green revolution”dwarfing genes *Rht-B1* and *Rht-D1* – knock-out mutations in *DELLA* proteins – into US and CIMMYT germplasm dramatically improved yields by increasing wheat harvest index and preventing lodging due to applied inorganic nitrogen fertilizer (Hedden, 2003). The effect of the *Rht1* genes is conditional on the environment and the quantity of assimilate produced by the variety, and has been associated with larger grain number but smaller grain size and weight (Borner et al., 1993). *Rht1* alleles disable plants’ ability to respond to giberellic acid (GA-insensitivity), which may have negative effects on coleoptile length and early plant vigor that can decrease yield in some environments (Rebetzke and Richards, 2000). Increase in seed number and grain yield seems to be related not to ear development but to the greater availability of assimilates during grain-fill with reduced biomass partitioned into stalks (Youssefian et al., 1992). Breeders generally select plants near some optimal height value, as too-short plants have a generally lower yield compared to semi-dwarfs characteristic of having only one *Rht* allele (Fischer and Quail, 1990). An increasing number of dwarfing genes in wheat have been fine-mapped, and many, though not most, have been cloned.

Genomic selection is restructuring modern wheat breeding programs. The ability to leverage data from past years to predict unobserved lines has tremendous potential to increase the rate of genetic gain. Beyond yield predictions, heading date and plant height predictions are valued by breeders, allowing them to exclude phenotypically extreme individuals without having to dedicate resources to planting and phenotyping in multiple environments. Standard GBLUP and rrBLUP models are optimized for highly polygenic traits like yield, but will underestimate QTL effect sizes and perform poorly with traits dominated by a smaller number of larger-effect QTL. Explicitly characterizing and taking into account large-effect QTL in traits where these QTL explain a substantial portion of additive genetic variation can increase prediction accuracy (Bernardo, 2014; Sarinelli et al., 2019). This may have even more promise in biparental populations where the number of segregating causal variants is much smaller. If traits are mostly controlled by a few major variants, models using markers for just those variants can be predictive and more cost-effective.

Here we set out to understand the genetic basis of plant growth traits in a biparental common wheat population. Parents were chosen to generate major additive genetic variation for plant height and heading date and to characterize novel QTL for these traits. Parent SS-MPV57 carries the large-effect earliness allele *Ppd-D1a* as well as the smaller-effect allele *Ppd-A1a*.*1*, but no known dwarfing genes. Parent LA95135 caries the major dwarfing allele *Rht-D1b* but no known earliness genes except the smaller-effect *Ppd-A1a*.*1* allele. This parental selection contrasts with typical mapping studies, where the two parents generally differ for the trait of interest. Here, due to our understanding of already characterized major variants, we developed a population with the goal of generating transgressive segregation for our target phenotypes. A high-density sequence-based linkage map was supplemented with single SNP assays for putative causal variants in order to map novel QTL and study marker-trait associations for known variants. Phenotypic variation in each environment was partitioned into components associated with mapped QTL and the polygenic background in order to assess the relative importance of identified QTL. Different models were tested for prediction of both traits to determine if a simple QTL model would be sufficient in the context of a breeding program. Finally, a longitudinal analysis of multiple measures of plant height over time was used to determine QTL effects over the course of plant growth in a field season.

## 2 Materials and Methods

### Population Development

Soft-red winter wheat lines developed by southeastern public-sector breeding programs were screened for alleles at known plant height and heading date variants using Kompetitive Allele-Specific PCR (KASP) markers. Louisiana State University forage cultivar LA95135 (CL-850643/PIONEER-2548//COKER-9877/3/FL-302/COKER-762) was chosen as a parent lacking major early-flowering alleles at the *Ppd-D1* or *Vrn-1* loci, but with a mid-season heading date when grown in North Carolina. Cultivar SS-MVP57 (FFR555W/3/VA89-22-52/TYLER //REDCOAT*2/GAINES) developed at Virginia Polytechnic Institute and State University displayed semi-dwarf stature but lacked dwarfing alleles at the *Rht1* loci. SS-MPV57 carries the *Ppd-D1a* allele conferring photoperiod insensitivity, and LA95135 has the *Rht-D1b* allele conferring semi-dwarfism. Parent lines were crossed, and F1 plants were selfed to generate an F_2_ population (hereafter referred to as the LM population). The F_2_ and later generations were inbred via the single-seed descent method until the F_5_ generation, producing 358 F_5_-derived recombinant inbred lines (RILs).

### Phenotyping

During the winter of 2016-2017, an experiment was conducted in the greenhouse to evaluate heading date. Imbibed seeds from each RIL were placed in a cold chamber kept at 4*°*C for 8 weeks and were transplanted into plastic cones (volume 0.7L, 6.5 cm in diameter and 25 cm depth) containing soil mix. Plants were grown in a completely randomized design with four replications in a greenhouse set at 16 hr photoperiod and 20*°*C /15*°*C (day/night) temperature.

To evaluate the impact of vernalization on the genetic architecture of heading date and on effects of individual QTL, the greenhouse experiment was repeated with a low-vernalization treatment in the winter of 2017-2018. This experiment was performed as above, except that imbibed seeds were placed in the cold chamber for only four weeks prior to transplanting. In addition, the LM RIL population was evaluated in the field at Raleigh, NC and Kinston, NC during the 2017-2018 season, and in Raleigh, Kinston, and Plains, GA in the 2018-2019 season, sown in the fall at the locally recommended times for commercial winter wheat production. The 358 RILs were grown using an augmented set within replications design to facilitate planting of this large population. RIL experiments consisted of two fully replicated blocks of all 358 lines organized into five sets of 71 or 72 RILs. The order of the sets within each replication and the order of the RILs within each set were randomized at each location. Three parental checks were planted at the beginning of each set of RILs, along with four or five additional parental checks randomized within each set.

Plots consisted of 1-m rows spaced 30 cm apart. Adult plant height was measured as the distance from the ground to the top of the spikes of a sample of tillers from the center of each row, excluding the awns. Heading date was measured as the day on which approximately half of the heads in each row had fully emerged from the flag leaf, typically a few days prior to anthesis. To study plant development over time, three measures of plant height were collected for each row plot in Raleigh in 2019, with two to four blocks measured roughly every ten days starting on March 29th and ending on April 25 (when most plants had fully headed). In this case, plant height at each time point was calculated as the mean of the height of three randomly chosen primary tillers from the ground to the base of the apical leaf sheath (Fig. 2). All measurements were collected on an android tablet with the Fieldbook app (Rife and Poland, 2014).

**Figure 2:**
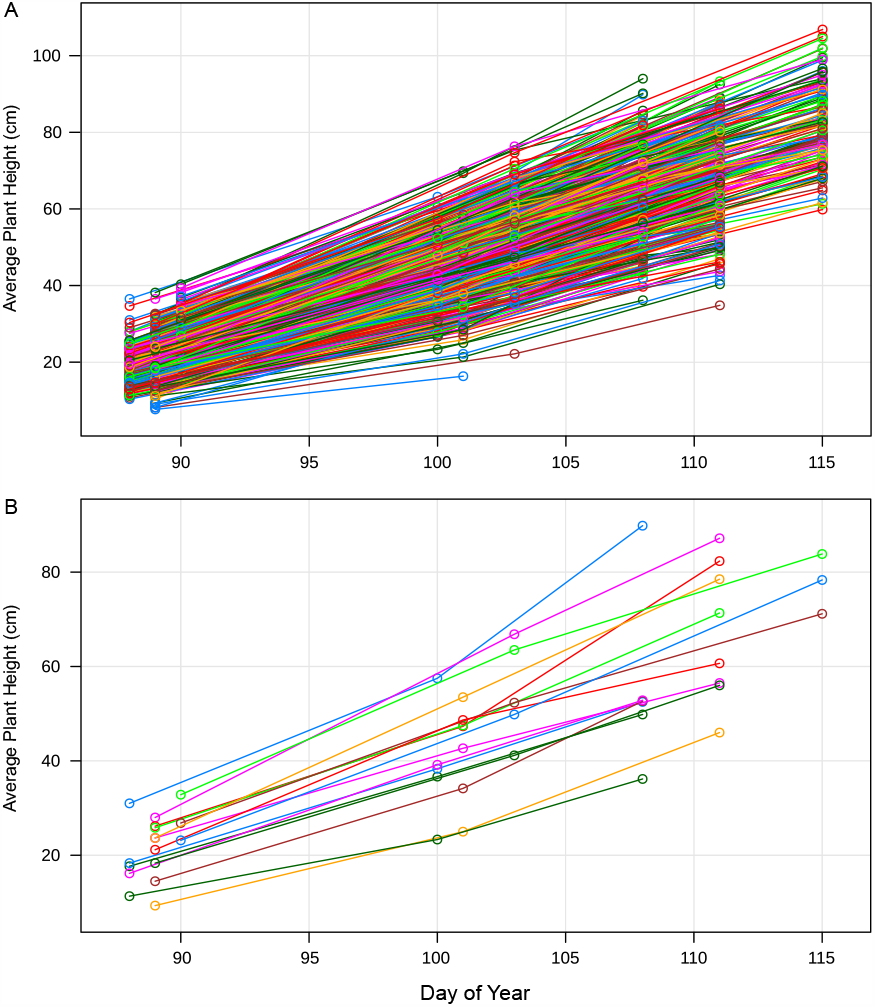
Plant growth over time. For each 1-m row plot (differently colored line), a total of three plant height values was collected in Raleigh in 2019. All plots are shown (A), as well as a random subset to better visualize plant growth (B). Mean plant growth follows a roughly linear pattern corresponding to the date collected, with different slopes and intercepts for each plot.

### Analysis of Phenotypes

For the greenhouse experiments where plants had been completely randomized within greenhouses, genotype values for RILs were calculated as the mean of the four replications of each line. For field experiments, best linear unbiased estimates (BLUEs) were calculated adjusting for these spatial effects. The software ASReml-R (Butler et al., 2017) was used to calculate BLUEs with an AR1xAR1 correlated residuals model:

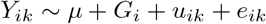

Where *Y*_*ik*_ is the observed phenotype for an individual row plot, *µ* is the intercept, *G*_*i*_ is the fixed effect of genotype *i, u*_*ik*_ is the unit or “nugget”random residual effect for each observation *k* representing the component of the variance due to observation or measurement instead of spatial correlation, drawn from a distribution 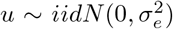, and *e*_*ik*_ is the spatially-correlated residual drawn from the distribution 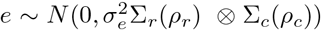, whose variance is the direct product of an *r × r* auto-correlation matrix Σ_*r*_(*ρ*_*r*_) representing autoregressive correlations in the row direction and *c × c* correlation matrix Σ_*c*_(*ρ*_*c*_) representing autoregressive correlations in the column direction. For all environments and phenotypes, a full model with autocorrelated columns and rows was found to have a lower BIC and higher log likelihood than models with just the column autocorrelation or no spatial correction. BLUEs were calculated as the sum of the genotype effect and the intercept for each phenotype in each environment.

### Genotyping and Linkage Map Construction

Tissue was collected from the F_5_ greenhouse experiment, and seeds of the four F_5:6_ plants from each line were bulked. Genotyping by sequencing (GBS; (Elshire et al., 2011)) was performed according to Poland *et al*. 2012 (Poland et al., 2012), with ninety-six individual samples barcoded, pooled into a single library, and sequenced on an Illumina HiSeq 2500. Tassel5GBSv2 pipeline version 5.2.35 (Glaubitz et al., 2014) was used to align raw reads to the International Wheat Genome Sequencing Consortium (IWGSC) RefSeqv1.0 assembly (https://wheat-urgi.versailles.inra.fr/Seq-Repository) using Burrows-Wheeler aligner (BWA) version 0.7.12 to call SNPs (Li et al., 2009). SNPs were filtered to retain samples with ≤20 percent missing data, ≥ 30 percent minor allele frequency and ≤10 percent of heterozygous calls per marker.

KASP markers taken from the literature or designed from exome capture data of the parents (triticeaetoolbox.org/wheat; Additional file 1: Table S1) were added to the GBS SNP data for chromosome regions with low marker density and for causal variants segregating in the population. Filtered SNPs were separated into chromosomes and ordered via alignment to the reference genome, and a custom script was run to filter out genotyping errors that would result in a false double recombination due to under or mis-calling of heterozygotes. The R package ASMap was used to construct the maps as an F_5_ RIL population (Taylor and Butler, 2017).

### QTL Analysis

QTL mapping was performed in the R package r/QTL (Broman et al., 2003). Composite interval mapping was used for initial QTL identification, and intervals were narrowed using a multiple QTL model (MQM) as implemented in the refineqtl function. The addqtl function was used to identify additional QTL using identified QTL as covariates. Empirical significance thresholds for a genomewide *α* = 0.05 were determined using 1000 permutations for each trait. QTL effects were estimated for significant QTL in each environment based on the estimated MQM positions using the fitqtl functions, which fits a multiple regression where for genotype values *Y*_*i*_ for each individual *i*, and 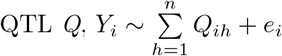.

Major variants *Ppd-D1* and *Rht-D1* alter functions of core genes in the flowering time and giberellic-acid response pathways, respectively, suggesting that their presence may alter the effects of other variants impacting those pathways. Taking advantage of the large number of lines in the LM population, two sub-populations of roughly 160 lines – each divided by genotype at the major-effect QTL – were created for mapping of each phenotype. Lines called as heterozygous for the major-effect QTL were excluded. The QTL mapping analysis was repeated for both of the sub-populations. Identified QTL interactions discovered this way were validated by modifying the above fitqtl model with a main effect for the identified QTL and an effect for its interaction with the major classifying QTL.

Variance analysis was performed in the R package lme4qtl, which allows for the fitting of random effects with supplied covariance matrices (Ziyatdinov et al., 2018). For known variants for which KASP marker genotypes of the causal polymorphisms were available, the genotypes were used directly, and for novel QTL genotype probabilities from the refineqtl object were used. For testing QTL, alleles were encoded in terms of the allele dosage of the LA95135 allele (0, 1, 2) without estimating a dominance effect.

While BLUEs estimated using the correlated errors model were used for QTL mapping, to estimate the relative importance of identified QTL in determining total phenotypic variation at the level of individual plots models were re-fit in each environment using the unadjusted phenotypes as the response. For each environment and phenotype, QTL effects and variance components for each the additive and non-additive effects of genotypes were specified with the mixed model:

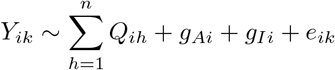

Where for each phenotype *Y* of genotype *i* in row plot *k*, fixed effects for each QTL *h* were fit as regressions of allele dosage on phenotypes. *g*_*Ai*_ represents the random additive effect of genotype *i* with a variance specified by the realized relationship matrix 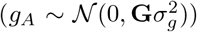, calculated using the A.mat function in the R package rrBLUP from the scaled GBS marker matrix (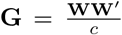, where **W** is the scaled marker matrix calculated as *W*_*ik*_ = *X*_*ik*_ + 1 − 2*p*_*k*_ from the frequency of the 1 allele at marker *k* (*p*_*k*_) and the marker matrix *X*_*ik*_. *c* is a normalization value calculated as *c* = 2Σ_*k*_*p*_*k*_(1 − *p*_*k*_)) (Endelman, 2011). *g*_*Ii*_ represents the non-additive random effect of genotype *i* with an independent variance 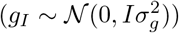.

A modified method from Nakagawa and Schielzeth 2013 (Nakagawa and Schielzeth, 2013) was used to estimate variances associated with QTL and variance components from the specified model. Estimated coefficients for fixed effects are multiplied by the value of that effect (in this case, the allele dosage), and the variance of these values is taken as the variance associated with that fixed effect. This *R*^2^-like estimator for mixed models is defined as:

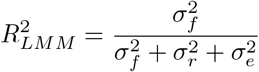

Where 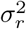 is the variance of the random effects and 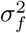 is a variance of independent fixed effects calculated as 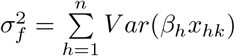 for coefficients and effects *h* and observations *k*. For both traits, all QTL were mapped to separate chromosomes, satisfying the assumption of independence. Using this approach, narrow-sense heritability in this population with *n* QTL in *i* individuals in *k* head rows is calculated as:

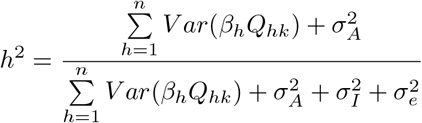

Where we calculate per-observation QTL effects as the allele dosage of QTL *h* in plot *k* (*Q*_*hk*_) times the estimated coefficient of each QTL (*β*_*h*_), and the phenotypic variance associated with that QTL as the variance of these estimates. The total variance associated with all QTL is taken as the sum of these individual QTL variances, as all mapped QTL are located on separate chromosomes and are independent of one another. 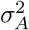 is the variance component associated with the random *g*_*A*_ genotype term fit with the relationship matrix, and 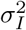 as the variance component associated with the random independent *g*_*I*_ genotype term, which represents some combination of epistatic effects, lack of linkage between observed markers and underlying causal variants, and deviation of the estimated genotype values from the true genotype values. Constructing the model in this way, we estimate the proportion of additive genetic variation associated with a QTL *h* (*p*_*A*_) as:

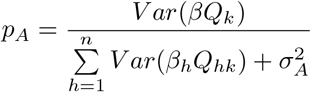

Where *p*_*A*_ is taken as the variance of the product of an estimated QTL effect by the allele dosage of that QTL in *k* rows, over the total additive genetic variation.

For investigating the effect of QTL on plant height variation over time, individual slopes of plant height over time measured multiple times for each row were calculated with a fixed intercept and a random intercept and time slope for each head row. The model was used to estimate plant height values for each row every day over the course of the month data was collected. A linear model fitting all relevant plant height and heading date QTL on plant height on every day was fit, and the partial *R*^2^ values of each QTL calculated for each day were used to estimate the relative importance of each QTL at each time point.

### Prediction of Phenotypes

Different prediction models were assessed to identify an optimal model for heading date and adult plant height. All models except for the simple QTL multiple regression model were fit in the R package BGLR (Pérez and De Los Campos, 2014), which allows for flexible fitting of a variety of Bayesian and mixed effects models. A GBLUP model was fit solving the equation *y* ∼ *µ* + *u* + *e* for *u. y* is a vector of BLUEs across environments for all RILs, with unobserved RILs assigned missing values, and *u* is a vector of random genotype effects with a variance 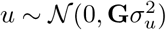, where **G** is the realized relationship matrix calculated previously from GBS markers.

A simple multiple-regression QTL model based on identified QTL was fit solving the equation 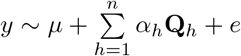 for *n* QTL, where **Q**_*h*_ encodes the LA95135 allele dosage for each QTL *h* in each individual, and *α*_*h*_ is the allele effect of QTL *h*.

A combined model was also fit specifying both a multiple-regression fixed-effects component for QTL effects, and random effects for each genotype constrained by the additive relationship matrix 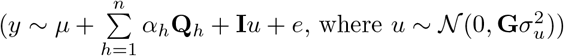.

BayesB and Bayesian LASSO models were both fit with the general model *y* ∼ *µ* + **X***u* + *e*, where **X** is a design matrix of markers coded by allele dosage of the LA95135 allele, and *u* is a vector of random marker effects. In the BayesB model, a certain proportion of markers given by the prior probability 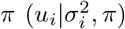 are assumed to have an effect size of 0, with the remainder having effects following a scaled-t distribution (Pérez and De Los Campos, 2014). In the Bayesian LASSO, marker effects were estimated with a double exponential prior distribution that assumes a greater frequency of both larger marker effects and marker effects closer to zero than a normal distribution (Pérez and De Los Campos, 2014). In both models, estimated genotype values are calculated as the sum of marker effects 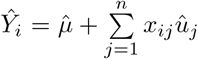, where *x*_*ij*_ is the allele dosage of marker *j* in individual *i*, and 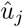 is the estimated marker effect.

A five-fold cross validation approach was used to compare the five models. RILs were randomly assigned to one of five folds, and genotype values from each environment from lines in four of the folds were used to predict the values of lines in the fifth fold, repeating for each fold in each environment for each model. Within each fold, QTL detection was re-performed as described in the QTL analysis section to identify the QTL used in the QTL regression and combined QTL and GBLUP model. This process was then repeated 40 times to get distributions of prediction abilities, calculated as the Pearson’s correlation between predicted and observed genotype values across all five folds.

## Results

### Genetic Map Construction

After filtering, 5691 markers were assigned to 21 linkage groups representing 21 wheat chromosomes. Average chromosome map length was 208.3 cM, with a maximum individual chromosome length of 319.1 cM for chromosome 3B. Marker density on the D genome tended to be much lower than marker densities on the A and B genomes, as expected given the much lower D genome diversity in hexaploid wheat (Akhunov et al., 2010).

### Population Characterization

Generally, wheat cultivars’ flowering habits are described by their genotypes at major heading date loci, but SS-MPV57 flowered later than LA95135 in all locations despite carrying the major earliness allele *Ppd-D1a* (Table 1). The difference in heading date was especially pronounced in the low-vernalization treatments both in the greenhouse (GH 2018) and in the field (Plains 2019), where SS-MPV57 flowered five and six days later, respectively, than LA95135. A similar pattern was observed for plant height: although LA95135 was the only parent genotyped for a major dwarfing allele (*Rht-D1b*), SS-MPV57 was substantially shorter in all locations (Table 1). For heading date in all locations, the mean genotype value of the RILs was approximately the mid-parent value. For plant height in Raleigh 2018 and Kinston 2018, the mean genotype value of the RILs was closer to the SS-MPV57 parent than the mid-parent value. The ranges of genotype values in Raleigh and Kinston were similar, but the range in heading date in Plains 2019 (26 days) was much larger. This is likely a result of the warmer winter temperatures at that site, delaying heading of lines with a greater vernalization requirement.

**Table 1:**
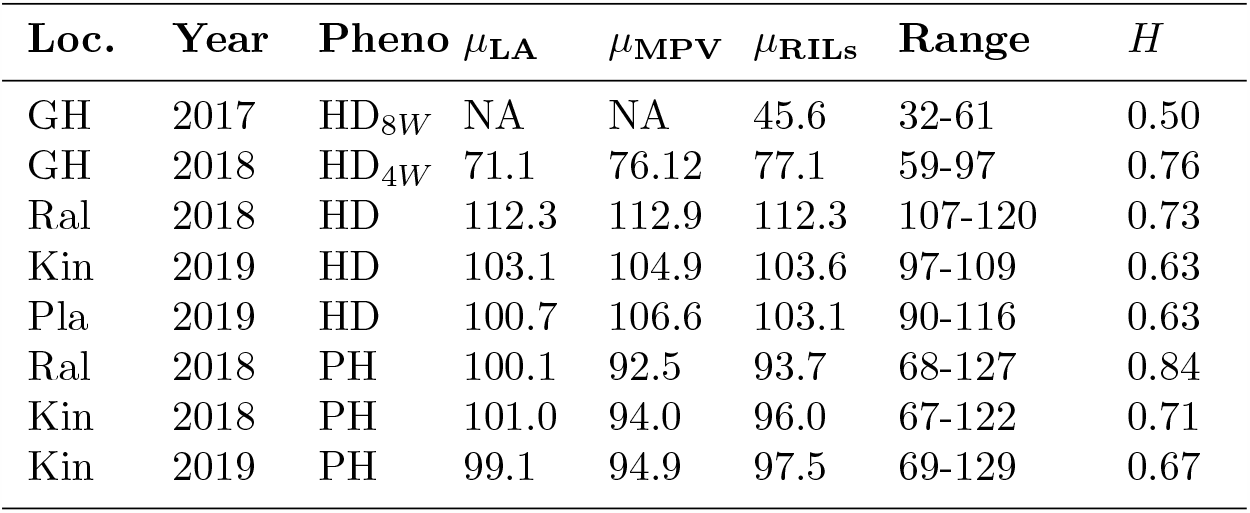
Population characteristics. Means and ranges of estimated genotype values for all RILs, as well as parental values and plot-basis heritabilities (H), for site-year-phenotype combinations. Heading date for the GH experiments is recorded as days since transplanting (four weeks (HD_4_*W*) or eight weeks (HD_8_*W*) after vernalization), and for the field experiments time as day of year (DOY).

### Known Variants and Novel QTL Impact Plant Growth

Genetic variation in quantitative traits like those measured in this study may result from the segregation of an unquantifiable number of small-effect QTL. Despite this, for both heading date and adult plant height the vast majority of additive genetic variation was associated with a small number of major QTL, some of which have been previously described and some of which are novel.

### Heading Date

The RIL population was developed with the expectation that the major photoperiod-insensitive allele *Ppd-D1a* inherited from SS-MPV57 would segregate, and that potential novel early-flowering QTL from LA95135 could be mapped. Two preliminary greenhouse experiments were conducted to investigate the effect of vernalization treatments on heading date genetic architecture, with imbibed seeds given only four weeks of vernalization in the first experiment and a full eight weeks in the second. In addition, heading date notes were collected in three separate field experiments in Raleigh in 2018, and Kinston and Plains, GA in 2019. QTL were declared significant at *α* = .05 based on 1000 permutations of the scanone function, but for all phenotypes significance values were near a LOD of 3.5. Together, *Ppd-D1, Rht-D1*, and four early-flowering alleles inherited from LA95135 were associated with differences in heading date in this experiment (Table 2, Table 3).

**Table 2:** Significant heading date QTL for four and eight week vernalization greenhouse experiments. The chromosome on which each QTL is found is indicated in the QTL name. For each QTL, the average difference in phenotype between two RILs homozygous for alternate alleles is given as twice the estimated allele effect of the LA95135 allele (2*α*), along with proportion of additive variation associated with each QTL (**p**_**A**_). The most significant markers for each QTL with a proposed candidate gene was a KASP marker associated with a previously identified causal polymorphism affecting that gene. Physical positions are given based on mapping of GBS markers to the IWGSC RefSeqv1.0 assembly.

| Treatment | QTL Name | Candidate Gene | Peak Marker | Position CI | LOD | $2\alpha$ (days) | $p_A$ |
| --- | --- | --- | --- | --- | --- | --- | --- |
| 4 Wk | *Qncb.HD-2D* | *Ppd-D1* | Ppd-D1 | 32-43 Mb | 11.5 | 5.2 | 0.15 |
| 4 Wk | *Qncb.HD-3A* | *FT2* | FT2 | 118-478 Mb | 21.2 | 6.5 | 0.25 |
| 4 Wk | *Qncb.HD-4D* | Rht-D1 | Rht-D1 | 0-352 Mb | 22.2 | 7.2 | 0.39 |
| 4 Wk | *Qncb.HD-5A* | NA | S5A_395681218 | 46-438 Mb | 5.65 | 3.2 | 0.05 |
| 4 Wk | *Qncb.HD-5B* | NA | S5B_462554252 | 427-523 Mb | 4.63 | 3.6 | 0.08 |
| 8 Wk | *Qncb.HD-2D* | *Ppd-D1* | Ppd-D1 | 33-44 Mb | 15.9 | 3.9 | 0.60 |
| 8 Wk | *Qncb.HD-3A* | *FT2* | FT2 | 71-435 Mb | 6.67 | 2.3 | 0.22 |
| 8 Wk | *Qncb.HD-5B* | NA | S5B_518684640 | 511-537 Mb | 5.17 | 2.1 | 0.14 |

**Table 3:**
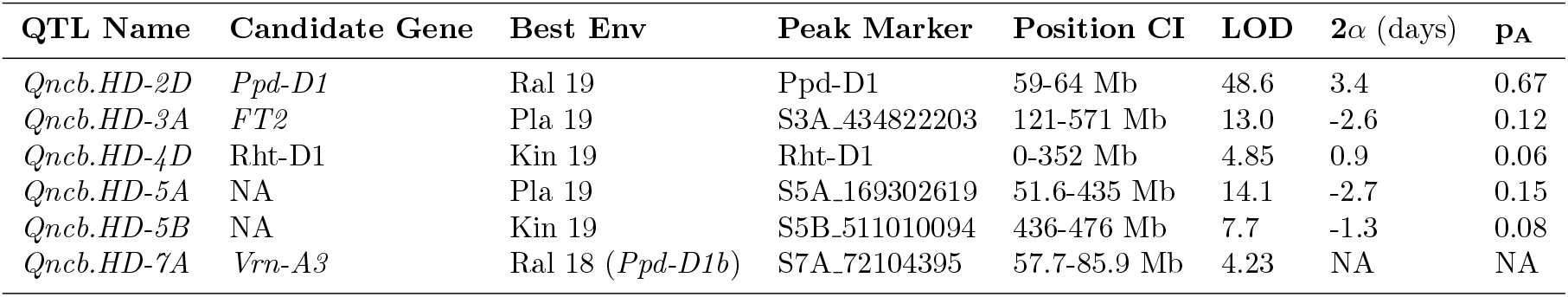
Significant heading date QTL information from best environment. For each QTL, information from the experiment where that QTL had the largest estimated effect (Best Env) is given. The average difference in phenotype between two RILs homozygous for alternative alleles at each QTL is given as twice the estimated allele effect of the LA95135 allele (2*α*), along with proportion of additive variation associated with each QTL (**p**_**A**_). *Vrn-A3* only has a significant effect within the half of the population homozygous *Ppd-D1b*.

| QTL Name | Candidate Gene | Best Env | Peak Marker | Position CI | LOD | $2\alpha$ (days) | $p_A$ |
| --- | --- | --- | --- | --- | --- | --- | --- |
| *Qncb.HD-2D* | *Ppd-D1* | Ral 19 | Ppd-D1 | 59-64 Mb | 48.6 | 3.4 | 0.67 |
| *Qncb.HD-3A* | *FT2* | Pla 19 | S3A_434822203 | 121-571 Mb | 13.0 | -2.6 | 0.12 |
| *Qncb.HD-4D* | *Rht-D1* | Kin 19 | Rht-D1 | 0-352 Mb | 4.85 | 0.9 | 0.06 |
| *Qncb.HD-5A* | NA | Pla 19 | S5A_169302619 | 51.6-435 Mb | 14.1 | -2.7 | 0.15 |
| *Qncb.HD-5B* | NA | Kin 19 | S5B_511010094 | 436-476 Mb | 7.7 | -1.3 | 0.08 |
| *Qncb.HD-7A* | *Vrn-A3* | Ral 18 (*Ppd-D1b*) | S7A_72104395 | 57.7-85.9 Mb | 4.23 | NA | NA |

A heading date QTL on chromosome 2D co-localized with a known major-effect variant altering expression of the D-genome copy of pseudo-response regulator gene *Photoperiod-1* (*Ppd-D1a*) (Beales et al., 2007). This was the major QTL mapped in this experiment, associated with by far the highest LOD score in both the field environments (Table 3) and the eight week greenhouse treatment (Table 2). In the four week treatment the relative importance of *Ppd-D1* was diminished, primarily as a result of changes in the effects of other QTL.

A QTL in the centromeric region of chromosome 3A is mapped with low physical resolution (*>* 400 Mb), owing to the low recombination rates found in these regions in wheat (Table 2, Table 3). Contained in this interval is *FT-A2*, an ortholog of *FT* previously described by Shaw *et al*. (Shaw et al., 2019) as an important component of the wheat flowering time pathway (Fig. 1). A KASP assay designed from a polymorphism within *FT-A2* was the peak marker for this QTL in both greenhouse experiments (Table 2), and had a much greater effect in the four week vernalization treatment than in the eight week treatment (Fig. 3). *Qncb*.*HD-3A* was also identified as significant in all field experiments, but with alternate peak markers in the long arm of chromosome 3A (Table 3).

**Figure 3:**
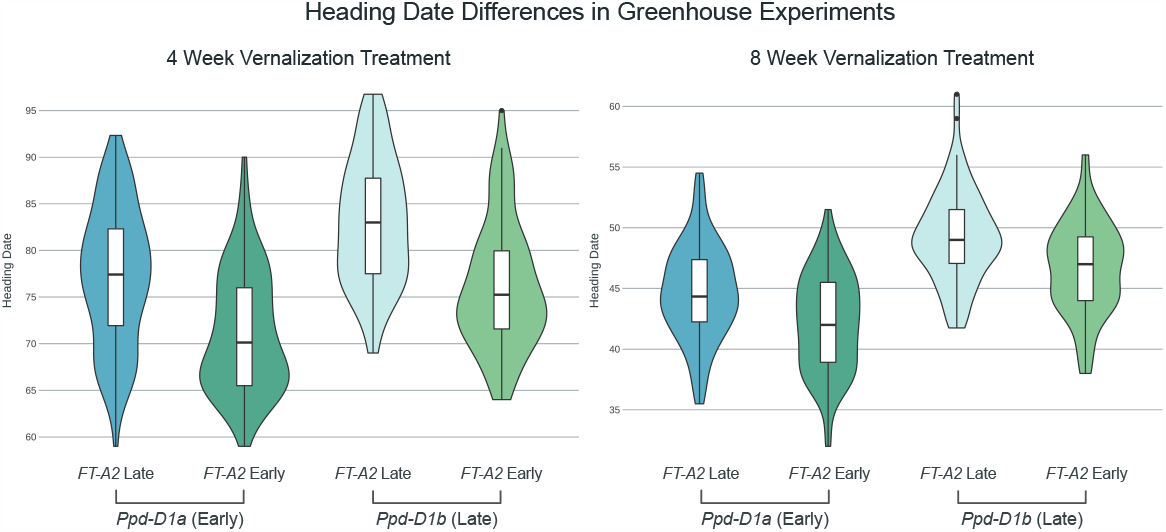
Effect of *Qncb*.*HD-3A* and *Ppd-D1* QTL on heading date in two different vernalization treatments. Density plots of BLUEs for heading date in two experiments, with RILs grouped by their genotype at *Ppd-D1* and a marker close to *FT-A2*. The allele effect of *Ppd-D1* is larger than that of *FT-A2* in the 8 week vernalization treatment (2.0 days versus 1.2), but the effect of the *FT-A2* marker is larger in the 4 week vernalization treatment (2.6 days vs 3.3 days).

In addition, two novel early-flowering alleles were identified on chromosomes 5A and 5B (Table 3). Both QTL are significant in all environments (Additional file 2: Table S3). *Qncb*.*HD-5A* is the third-most important QTL in most environments, but has an especially large effect in the Plains, GA field experiment. The QTL is also significant in the four week vernalization greenhouse treatment, but not the eight week treatment. The increased QTL effect in these two environments having shorter duration of cold temperature exposure suggests that *Qncb*.*HD-5A* may interact with genes involved in vernalization response. Due to its centromeric position, the confidence interval for the QTL contains 383 Mb of chromosome 5A (Table 3). Notably, despite the response of this QTL to vernalization treatment, this interval does not encompass the *Vrna-A1* locus.

*Qncb*.*HD-5B* is located in a more distal position on the long arm of the chromosome and was mapped to an interval of 61 Mb. This interval is proximal to *Vrn-B1*, excluding that locus as a candidate gene. Unlike *Qncb*.*HD-5A*, significance and effect sizes of *Qncb*.*HD-5B* are similar in both the four and eight week vernalization treatments (Table 2).

The major plant height QTL *Rht-D1* was also identified as having a pleiotropic effect on heading date in this population. In most environments the effect on heading date was minor, and not significant in the eight week greenhouse treatment or in Plains in 2019. However, in the four week greenhouse treatment *Rht-D1* was a highly significant QTL, with an average difference of over seven days between plants homozygous for wild type or semi-dwarf alleles (Table 2).

A benefit of large population sizes is the ability to subset the population by major-effect variant allele and perform QTL analyses on the sub-populations. In the case of the *Ppd-D1a* insensitive allele, constitutive over-expression of *Ppd-D1* may obscure effects of variation elsewhere in the flowering pathway. After dividing the population by *Ppd-D1* allelic class and performing QTL analyses on the sub-populations, an additional early-flowering allele from LA95135 was identified on the short arm of chromosome 7A only in a *Ppd-D1b* photoperiod-sensitive background, and only in the field experiments (Table 3). The confidence interval for this QTL contains the *Vrn-A3* locus. *Vrn3* in wheat was identified as an *FT* ortholog (*TaFT1*), and serves as the primary integrator of flowering time signal, being translocated from the leaves to the shoot apical meristem to initiate the transition to reproductive growth (Fig. 1) (Yan et al., 2006). A variant in the D-genome copy of this gene, *Vrn-D3a*, was identified by (Chen et al., 2010) as a determinant of flowering time in winter wheat. A deletion of a GATA box in the promoter region of *Vrn-A3* has been recently associated with delayed flowering time in tetraploid durum wheat (Nishimura et al., 2018), and an additional polymorphism linked to differences in heading date and spikelets-per-spike has also been identified in common wheat (Chen et al., 2020). Screening the population with a KASP marker developed around the GATA box deletion (Additional file 1: Table S1) reveals that the population segregates for the deletion, with SS-MPV57 contributing the late-flowering deletion allele. While *Qncb*.*HD*.*7A* has a relatively small additive effect, it strongly interacts with *Ppd-D1a* (Fig. 4). In a background containing the insensitive over-expression *Ppd-D1* allele, there is no difference in heading date between lines with and without the *Vrn-A3* promoter deletion. In a *Ppd-D1b* background, however, the GATA box deletion is associated with significantly delayed heading date of approximately one day (Fig. 4). In wheat, *Ppd1* acts to trigger expression of *Vrn3* through signaling intermediates (Fig. 1), thus an interaction between the two fits with our understanding of their placement in a common pathway. This promoter deletion is a strong candidate for the variant underlying the chromosome 7A heading date QTL.

**Figure 4:**
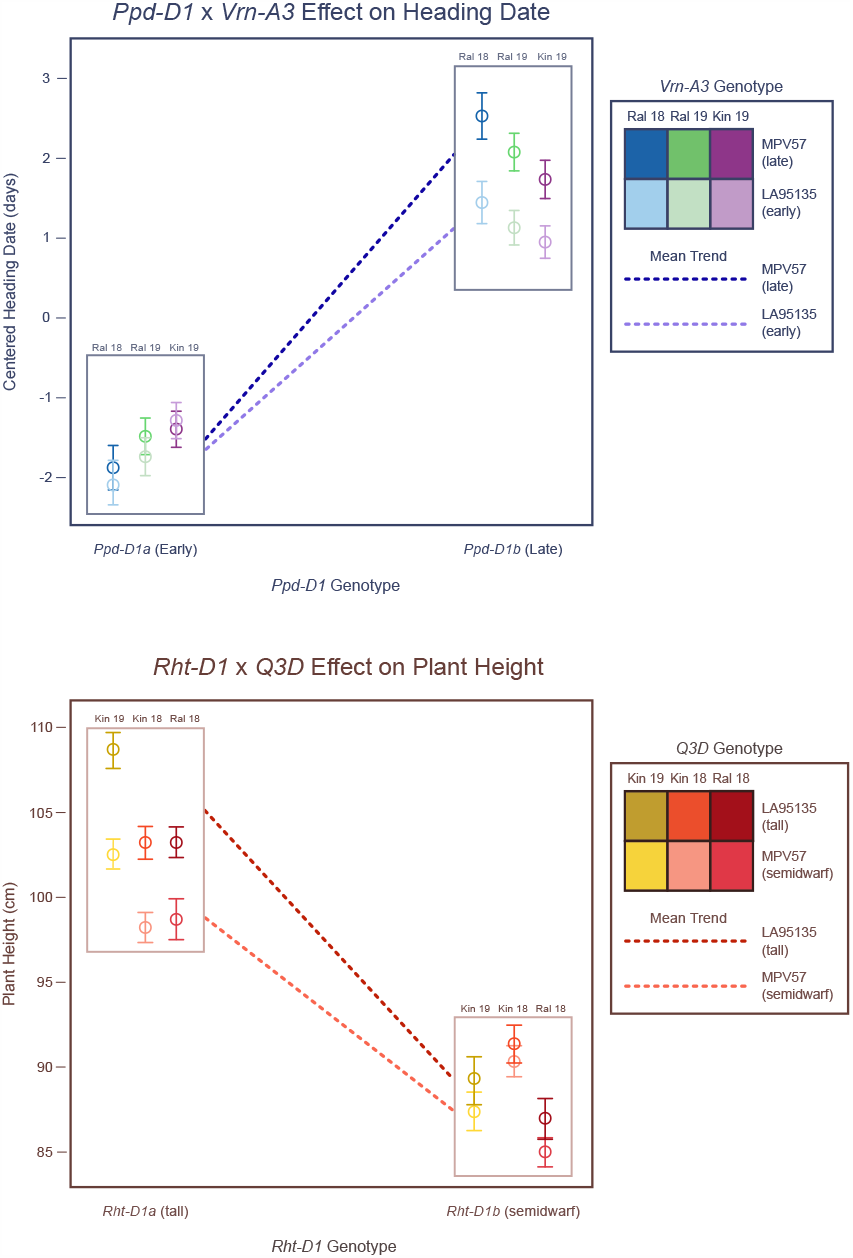
Major variants diminish effects of other QTL. *Vrn-A3* alters heading date in most environments, but only in a *Ppd-D1* sensitive background. The dwarfing effect of *Qncb*.*PH-3D* is greater in an *Rht-D1a* (tall) background.

### Adult Plant Height

Major QTL for plant height were initially mapped to chromosomes 4D, 2D, and 3D (Table 4). Using the MQM model, additional adult plant height QTL on chromosomes 1A, 2B, and 5B were also identified (Table 4). As expected, known variant *Rht-D1b* inherited from LA95135 was by far the largest-effect QTL across environments. Except for *Qncb*.*PH-5B*, all other reduced plant height alleles were inherited from SS-MPV57.

**Table 4:**
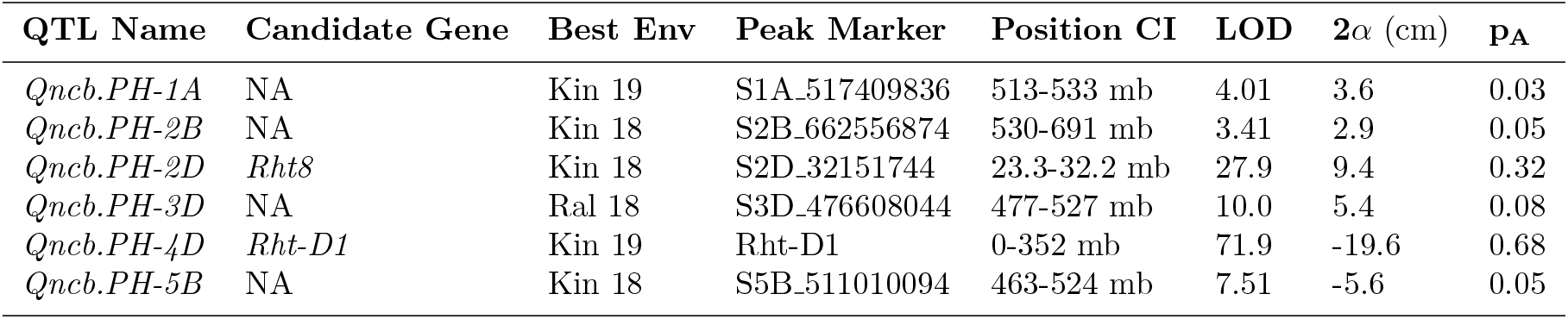
Significant plant height QTL information from best environment. For each QTL, information from the experiment where that QTL had the largest estimated effect (Best Env) is given. The average difference in phenotype between two RILs homozygous for each QTL is given as twice the estimated allele effect of the LA95135 allele (2*α*), along with proportion of additive variation associated with each QTL (**p**_**A**_). The confidence interval for *Rht8* is consistent with prior studies placing the QTL distal to *Ppd-D1*.

| QTL Name | Candidate Gene | Best Env | Peak Marker | Position CI | LOD | $2\alpha$ (cm) | $\mathbf{p_A}$ |
| --- | --- | --- | --- | --- | --- | --- | --- |
| *Qncb.PH-1A* | NA | Kin 19 | S1A_517409836 | 513-533 mb | 4.01 | 3.6 | 0.03 |
| *Qncb.PH-2B* | NA | Kin 18 | S2B_662556874 | 530-691 mb | 3.41 | 2.9 | 0.05 |
| *Qncb.PH-2D* | *Rht8* | Kin 18 | S2D_32151744 | 23.3-32.2 mb | 27.9 | 9.4 | 0.32 |
| *Qncb.PH-3D* | NA | Ral 18 | S3D_476608044 | 477-527 mb | 10.0 | 5.4 | 0.08 |
| *Qncb.PH-4D* | *Rht-D1* | Kin 19 | Rht-D1 | 0-352 mb | 71.9 | -19.6 | 0.68 |
| *Qncb.PH-5B* | NA | Kin 18 | S5B_511010094 | 463-524 mb | 7.51 | -5.6 | 0.05 |

The mapped position of the plant height QTL located on chromosome 2D is consistent with reported positions for *Rht8* (Gasperini et al., 2012). After the two major gibberellic-acid insensitive dwarfing genes *Rht-D1b* and *Rht-B1b*, the most commonly used gene is *Rht8*, which is tightly linked to *Ppd-D1* (Worland et al., 1998). In most environments, the marker most closely associated with *QPH*.*ncb-2D* is mapped closely distal to *Ppd-D1*. In an effort to tease apart the effects of photoperiod insensitivity and *Rht8* on plant height, we evaluated a terminal spike-compaction phenotype often associated with *Rht8* segregating in the LM population. This trait was rated in the field in Raleigh in 2019, and the major QTL was co-located with the plant height locus on the short arm of chromosome 2D (Additional file 1: Figure S1). We did not observe any significant interaction between *Rht8* and *Rht-D1*.

We identified *Qncb*.*PH-3D* as a novel plant height QTL, with a smaller effect than either *Rht-D1* or *Rht8* (Table 4). Despite the low marker density on chromosome 3D, *Qncb*.*PH-3D* was consistently localized to a 50-Mb interval on the long arm. *Rht-D1b* alters the function of an important component of the giberellic acid response pathway, so we may expect differential QTL effects in different *Rht-D1* backgrounds. We find that while *Qncb*.*PH-3D* was identified in all environments, the effect on plant height is much greater in a *Rht-D1a* (tall) background (Fig. 4). As SS-MPV57 is responsive to giberellic acid, the observed interaction between *Rht-D1* and *Qncb*.*PH-3D* will require further study, and may point to the identification of candidate genes for this QTL.

Three additional QTL (*Qncb*.*PH-1A, Qncb*.*PH-2B*, and *Qncb*.*PH-5B*) were also identified in one environment each, but when fit in the combined multiple QTL model all were significant with *p <* .001 in all environments (Additional file 2:Table S4).

### QTL with Major and Moderate Effects Explain Most of Additive Genetic Variation and Generate Transgressive Segregation

Within-field phenotypic variance was partitioned in order to assess the genetic architecture of plant growth traits in this population and the relative importance of different mapped QTL in explaining observed differences (Fig. 5). For both heading date and adult plant height, major effect QTL dominate additive genetic variation in most environments. Major-effect variant *Ppd-D1* was associated with a majority of additive genetic variation for heading date, except in the southern-most location of Plains, GA in 2019 (Fig. 5). In this environment, the polygenic additive genetic variation for heading date was similar to that associated with *Ppd-D1*. The modified architecture in a distinct environment suggests the presence of QTL with smaller effects conditional on photoperiod and vernalization signal. *FT2* and *Qncb*.*HD-5A* also increased in importance in the Plains 2019 environment, indicating that the effects of these moderate-effect QTL may also vary based on environmental conditions. Major-effect variant *Rht-D1* explained a majority of the additive genetic variation for plant height except in Kinston, NC in 2018 where *Rht8* explained a similarly sized proportion of variation (Fig. 5). The relative expression of these QTL in specific environments plays a large role in determining the observed variation both in genotype values and in phenotypes.

**Figure 5:**
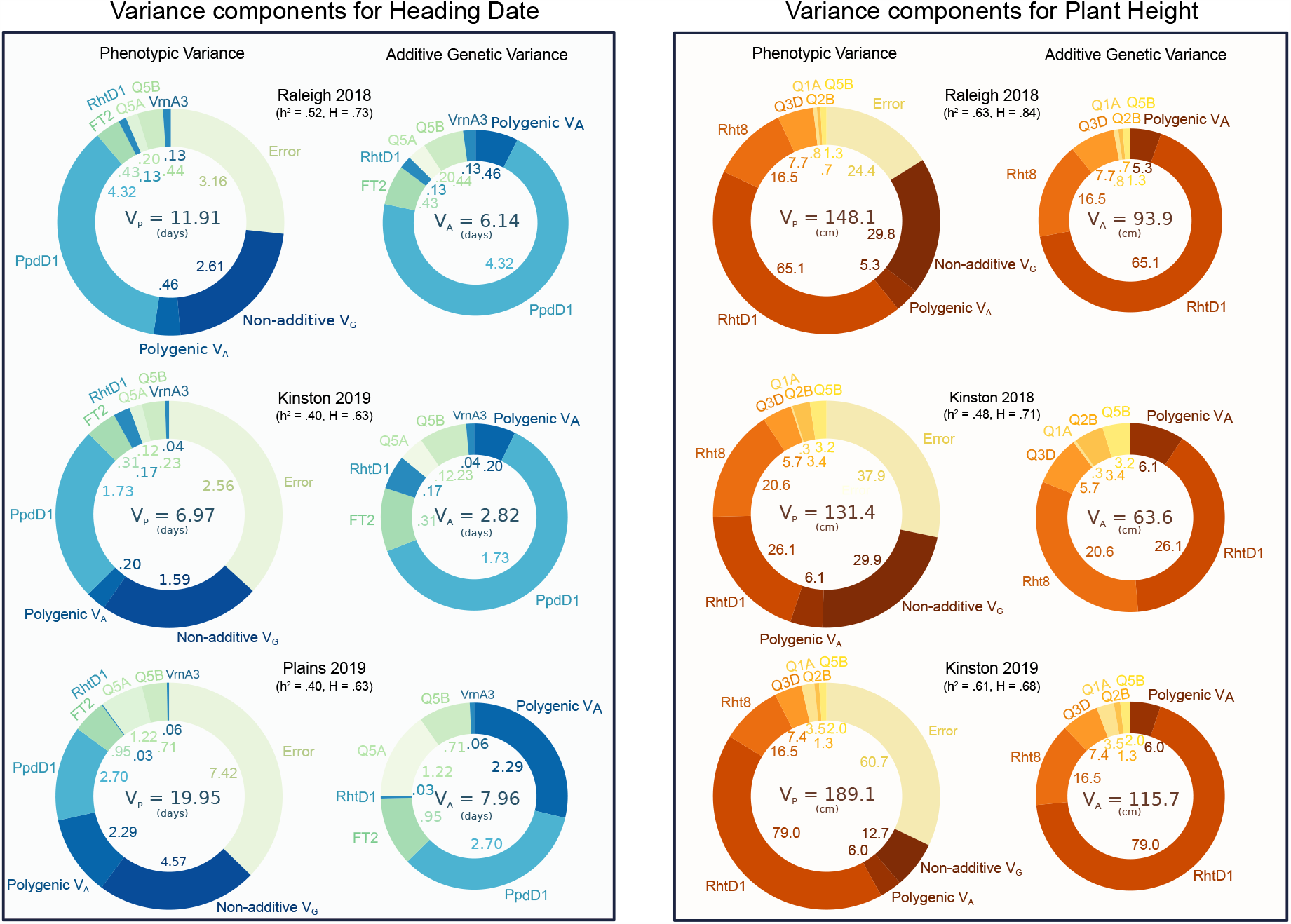
Variance associated with QTL and variance components for heading date and plant height in multiple environments. Non-additive genetic variation may be a result of epistatic interactions between QTL or mis-estimation of genotype values. *Ppd-D1* and *Rht-D1* dominate additive genetic variation for their respective phenotypes, but other mapped QTL explain a substantial portion of genetic variation. The scaling of total additive genetic variation is in large part due to the expression of *Ppd-D1* or *Rht-D1* effects.

A central question of this study is if variation in these plant growth traits is largely attributable to segregation of large-effect variants, or if identified variants are instead only contributing to largely polygenic differences in heading date and plant height. The plant height and heading date characters of the parental lines were found to be almost entirely determined by either major *Ppd-D1* and *Rht-D1* alleles, or cumulative effects of the stable, moderate-effect QTL identified in this study (Fig. 6). The transgressive segregation observed in this study, where both parents are phenotypically similar in terms of heading date and plant height, is being driven primarily by segregation of these major and moderate-effect QTL.

**Figure 6:**
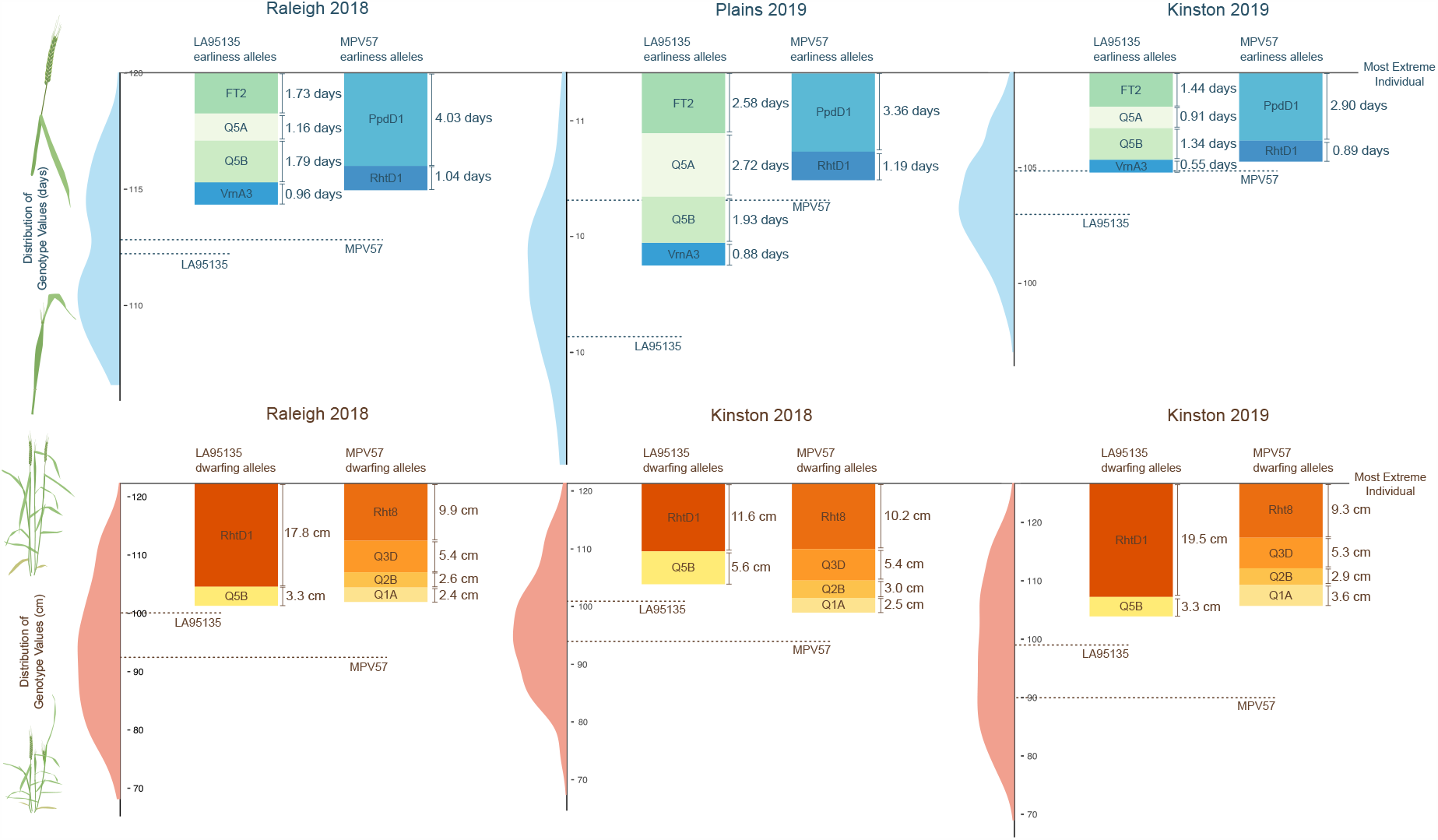
Heading date and plant height characters of parental lines are mostly determined by major QTL. For both heading date and plant height, the most phenotypically extreme individual was considered as the baseline for each environment and compared to both the distribution of genotype values and estimated QTL effects for the difference between two inbred lines (2*α*). Observed genotype values for the parental lines in each environment (dashed lines) are compared to the cumulative effects of their alleles.

For heading date, the effects of *Ppd-D1* and *Rht-D1* were mostly sufficient to explain the observed phenotypes of SS-MPV57, and the phenotypes of LA95135 were mostly explained by the QTL effects of earliness alleles inherited from that parent (Fig. 6). In Raleigh in 2018, *Ppd-D1* has the largest effect, visible in the apparent bimodal distribution of genotype values. Plants in this environment experienced the coolest winter temperatures and had the latest mean heading dates (Table 1). The differences between the two parents is greatest in Plains in 2019, where the effect of *Ppd-D1* is relatively reduced and larger effects are observed for earliness alleles inherited from LA95135. Plants in this environment experienced the warmest winter temperatures and had the greatest range in heading dates.

For plant height, the effect of *Rht-D1b* largely determines the semi-dwarf character of LA95135, along with some contribution from novel plant height QTL on chromosome 5B. The semi-dwarf character of parent SS-MPV57 is largely generated by known dwarfing QTL *Rht8* and the novel QTL on chromosome 3D, with some contribution from novel QTL on chromosomes 1A and 2B.

### QTL Models Out-Perform Genomic Selection for Oligogenic Traits

To assess the implications of the apparent oligogenic architecture of plant growth traits, a five-fold cross validation approach was used comparing a standard GBLUP model using genome-wide GBS markers to a simple multiple-regression QTL model based on previously estimated QTL effects (Table 5, Table 6).

**Table 5:** Prediction accuracies for heading date. Mean prediction abilities and their standard deviations estimated from 40 replications of five-fold cross validations using QTL regression, GBLUP, QTL fixed effects plus GBLUP, Bayes B, and Bayesian Lasso models.

| Model | Ral18 |  | Kin19 |  | Pla19 |  |
| --- | --- | --- | --- | --- | --- | --- |
| | $\mu$ | sd | $\mu$ | sd | $\mu$ | sd |
| QTL Regres. | 0.67 | 0.010 | 0.63 | 0.015 | 0.60 | 0.004 |
| GBLUP | 0.38 | 0.025 | 0.39 | 0.025 | 0.53 | 0.021 |
| QTL/GBLUP | 0.70 | 0.008 | 0.66 | 0.010 | 0.64 | 0.008 |
| Bayes B | 0.71 | 0.012 | 0.68 | 0.018 | 0.64 | 0.016 |
| Bayes Lasso | 0.63 | 0.018 | 0.58 | 0.024 | 0.63 | 0.022 |

**Table 6:** Prediction accuracies for plant height. Mean prediction abilities and their standard deviations estimated from 40 replications of five-fold cross validations using QTL regression, GBLUP, QTL fixed effects plus GBLUP, Bayes B, and Bayesian Lasso models.

| Model | Ral18 |  | Kin18 |  | Kin19 |  |
| --- | --- | --- | --- | --- | --- | --- |
| | $\mu$ | sd | $\mu$ | sd | $\mu$ | sd |
| QTL Regres. | 0.77 | 0.004 | 0.69 | 0.007 | 0.79 | 0.005 |
| GBLUP | 0.24 | 0.032 | 0.35 | 0.024 | 0.26 | 0.032 |
| QTL/GBLUP | 0.80 | 0.005 | 0.74 | 0.007 | 0.81 | 0.006 |
| Bayes B | 0.79 | 0.007 | 0.71 | 0.009 | 0.79 | 0.007 |
| Bayes Lasso | 0.67 | 0.015 | 0.50 | 0.030 | 0.70 | 0.014 |

Across all environments for both phenotypes, the simple QTL regression model is nearly as predictive as the top-performing model incorporating genome-wide marker information. The GBLUP model commonly used in applied wheat breeding is comparatively ineffective in predicting heading date and especially plant height within the biparental population. Incorporation of genomic relationship information into the QTL regression model only offers slight performance increases compared to the base model, suggesting the genomic relationships do not add much additional information. The Bayes B model, designed to allow for marker effects of zero, performs the best for heading date (Table 5). For plant height, the GBLUP model with QTL fixed effects is superior (Table 6). In general, the Bayesian Lasso model is superior to the GBLUP model but inferior to the other models, except for heading date in Plains in 2019 where the relative proportion of additive genetic variation associated with the polygenic background was the highest.

### Variation in plant growth is generated by major QTL

Plant height variation before maturity is caused in part by differences in development related to heading date variation, and thus may be controlled by QTL for both mature plant height and heading date. Multiple measures of plant height were collected from the RIL population planted in Raleigh during the 2019 field season, and a longitudinal model was used to estimate plant height over the measured time window. Identified heading date and adult plant height QTL were fit in a multiple regression model to estimate the proportion of phenotypic variation in plant height on a given day associated with each QTL. Variation in simulated genotype values were normalized by total QTL variation explained, and plotted over time to assess the relative importance of QTL in variation in plant height over time (Fig. 7).

**Figure 7:**
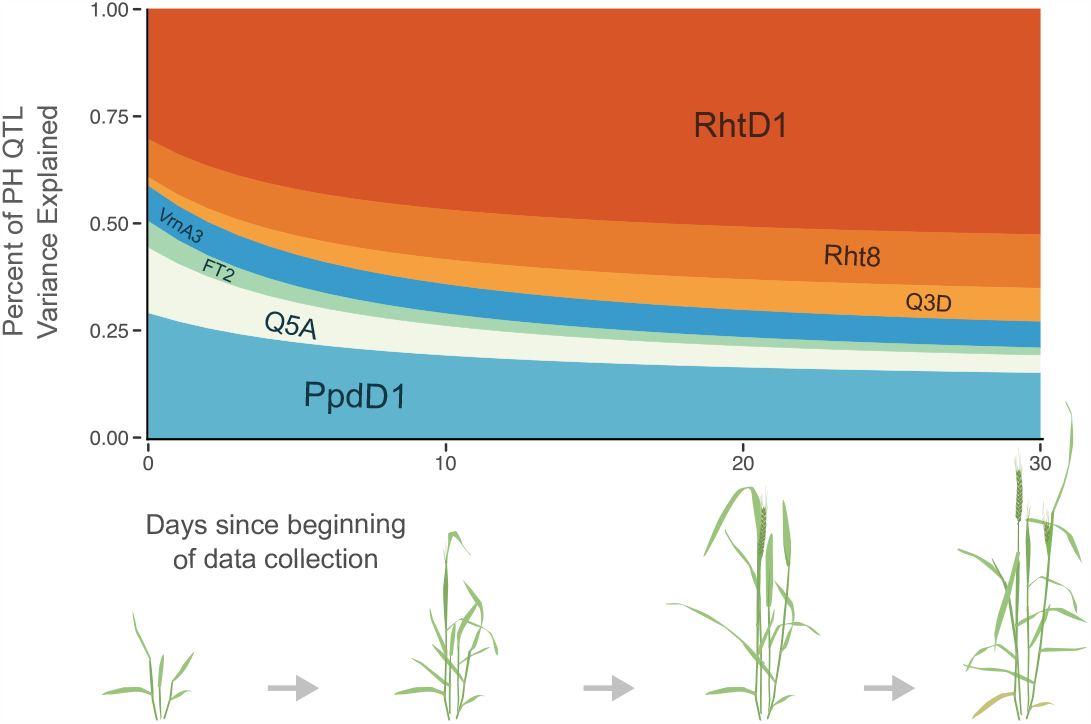
Relative importance of QTL for plant height over time. QTL associated with heading date (blue and green; *Ppd-D1, Qncb*.*HD-5A, FT2*, and *Vrn-A3*) explain over half of plant height variation associated with QTL at the beginning of data collection, but explain only approximately a quarter thirty days after data collecting began. The relative importance of plant height QTL (orange; *Rht-D1, Rht8*, and *Qncb*.*PH-3D*) increases over time.

As expected, the proportion of variation explained by the three adult plant height QTL (*Rht-D1, Rht8*, and *Qncb*.*PH-3D*) increases towards the end of the date range (March 29 to April 29, from near winter dormancy release to heading). Heading date QTL are more important than adult plant height QTL for early season plant height, when plants transition from vegetative to reproductive growth. The four heading date loci (*Ppd-D1, Qncb*.*HD-5A, FT-A2*, and *Vrn-A3*) continue to explain a large portion of variation in plant height as plants near heading, although their contribution diminishes as plants mature. Interestingly, the proportion of variation explained by QTL associated with *Vrn-A3* and *Rht8* were relatively consistent throughout development. *Rht-D1*, mapped as both a heading date and adult plant height QTL in this population, is associated with a large proportion of phenotypic variation throughout the date range.

## Discussion

### Unexplained Parental phenotypes result from novel QTL

Understanding the genetic basis of plant development is critical for understanding genetic variation for yield. In wheat, early flowering and plant height are understood to be largely determined by known large-effect variants; the mutations in the *DELLA* protein RHT1 (reduced height 1) on chromosomes 4B and 4D for plant height, and variation in vernalization-response *Vrn1* and photoperiod-response *Ppd1* genes for flowering time. Breeders generally select plants with plant heights and heading dates near some optimal values for their target environments, so that most cultivars have one of *Rht-B1b* or *Rht-D1b* but not both. In the southeastern US, most cultivars have some combination of early-flowering winter alleles of the *Vrn1* loci and one or more insensitive alleles of the *Ppd1* loci. Despite this, some cultivars with near-optimal values for heading date and plant height do not carry any known early flowering time or dwarfism alleles (for example, the two parents used in this study), and the relative importance of these major QTL versus other, smaller effect QTL in generating genetic variation for plant height and heading date is not known.

In other crop species such as maize, the majority of additive genetic variation in heading date and adult plant height is generated in a polygenic manner through the combination of many small-effect, unmapped QTL (Buckler et al., 2009; Peiffer et al., 2014). The importance of major-effect QTL in wheat (and other selfing species such as rice (Huang et al., 1996)) suggests that these traits may have a less polygenic basis in these species. Here we developed a biparental population by crossing cultivar LA95135, a cultivar with a normal flowering time but no early-flowering variants other than the weak photoperiod insensitive allele *Ppd-A1a*.*1*, to SS-MPV57, a cultivar with a normal height but no known *Rht1* variants. Within this population, additive genetic variation for plant growth phenotypes emerges from known major-effect QTL and multiple novel moderate-effect QTL, instead of primarily from polygenic background effects of many small-effect QTL.

We find one plant height QTL on chromosome 2D, mapped distal to *Ppd-D1*, that likely represents *Rht8*. Cultivars having the *Rht8* dwarfing allele are responsive to gibberrellic-acid (Korzun et al., 1998), and the gene has a smaller effect on plant height than reported effects of *Rht-B1* and *Rht-D1*, in agreement with the allele effects estimated in this study (Rebetzke et al., 2012). Additionally, we propose newly characterized variants in genes *FT2* and *Vrn-A3* as candidates underlying QTL on chromosomes 3A and 7A, respectively. Additional novel plant height QTL were mapped to chromosomes 3D, 1A, 2B, and 5B, and additional heading date QTL to chromosomes 5A and 5B. When considered jointly, the effects of these QTL and *Ppd-D1* and *Rht-D1* are mostly sufficient to explain the phenotypes of the parental lines. Our ability to identify these novel QTL despite their comparatively small effect size may be attributable to the large population size, twice that of many winter wheat RIL populations.

Combining these moderate-effect QTL can produce plants with a short enough height and an early enough heading date. Like other non-*Rht1* dwarfing genes, *Rht8* and *Qncb*.*PH-3D* do not confer GA-insensitivity to SS-MPV57 (data not shown). A major limitation of the GA-insensitive *Rht1* genes is a reduced coleoptile length, which can lead to poor emergence and weak competition. While lines carrying *Rht8* alone are often too tall, semi-dwarf lines like SS-MPV57 produced by stacking *Rht8* with *Qncb*.*PH-3D* may perform better than *Rht1* semi-dwarfs in certain environments (Worland et al., 1998). The insignificant effect on plant height of *Qncb*.*PH-3D* in an *Rht-D1b* (semidwarf) background also reduces the potential of producing transgressive segregants that are too short from crosses between *Rht-D1b* and *Qncb*.*PH-3D* -dwarf cultivars. The position of *Qncb*.*PH-3D* distally on the long arm facilitates its fine-mapping, and identification of a predictive marker or the underlying causal polymorphisms will facilitate marker-assisted selection of this QTL in developing GA-sensitive semi-dwarf cultivars.

The use of major *Ppd1* and *Vrn1* variants in cultivar development also has associated drawbacks. In both cases, the early-heading character is a result of the plant losing its ability to receive signal from its environment – in wild-type photoperiod-sensitive genotypes, plants use information about changing night lengths to flower at an appropriate time, whereas photoperiod-insensitivity activates this pathway constitutively. Losing the ability to respond to environmental cues may incur yield penalties in some situations. For example, autumn sown wheat lines with little or no vernalization requirement that are insensitive to photoperiod are susceptible to late spring freeze. However, requiring a long period of cold to potentiate flowering in environments with warm winters can result in delayed heading, even in lines having photoperiod insensitive alleles. This effect was observed in this study with the proportionally decreased effect of *Ppd-D1* in the Plains environment, which is farther south than the other locations and has shorter nights during the wheat growing season. A set of early flowering QTL with different environmental triggers or of more moderate effects, like those mapped here, provide breeders with additional tools to develop appropriate cultivars for various target environments. Fine-mapping and characterization of *Qncb*.*HD-3A, Qncb*.*HD-5A*, and *Qncb*.*HD-5B* will expand the flowering time toolbox for wheat breeders.

### The oligogenic trait architecture of plant growth traits in wheat

In wheat, genetic variation in yield is dependent on yield components (e.g. kernel weight, kernel number per spike, spikelets per spike) that can be influenced by disease resistance, plant height, and heading date, among other traits. While yield variation itself is complex, this complexity may arise through a combination of variation in other traits which may not necessarily have a polygenic architecture. We observe only a small fraction of the total genetic variation for heading date and plant height in this population associated with lines’ polygenic background. Even when not considering major-effect alleles *Rht-D1* and *Ppd-D1*, the remaining moderate-effect QTL explain more than twice the additive genetic variation as the polygenic background. While it is impossible to extend the results of this biparental study to wheat generally, the variation in heading date and plant height observed in this population is similar to the range of values observed in preliminary yield trials in breeding populations. It is likely the case that, while the particular variants differ from population to population, that the genetic architecture of plant height and heading date are similar across breeding populations in wheat.

### Challenges and opportunities for genotype-based prediction of plant growth traits

The genetic architecture of plant growth has important implications for modern wheat breeding programs. Yield is the primary target of wheat breeders, and standard genomic selection models perform well for this trait in southeast U.S. wheat breeding programs (Sarinelli et al., 2019; Ward et al., 2019). Standard models shrink estimated effects of large-effect variants closer to zero, which will reduce accuracy of models for traits mostly conditioned by relatively few large-effect variants (Bernardo, 2014). Given the effects of heading date and plant height variation on generating yield variation in wheat, if a handful of major QTL dominate these traits they may also have large effects on yield, complicating assumptions of these models. At the same time, heading date and plant height are themselves traits of interests to breeders, who screen biparental populations to remove transgressive segregants for these traits.

We show that the majority of additive genetic variation for heading date and plant height is controlled by large-effect QTL, such that a simple QTL model is sufficient for accurate prediction of phenotypes. In this case given marker information for major and moderate-effect QTL and a genotyped training population, a simple QTL model is likely to be effective for eliminating transgressive segregants for heading date and plant height. This model has the added benefit of being much cheaper than genomic selection if markers for polymorphisms linked to variants are available. Instead of genotyping a population of a set size with genome-wide markers, making predictions with genomic selection models, and then removing transgressive segregants for plant height and heading date, breeders can instead screen larger populations initially with simple makers for major QTL, and focus genotyping resources on lines predicted to be near optimal values for those secondary phenotypes. While QTL mapping was necessary to identify many important QTL for prediction in the QTL regression model in this population, this population was constructed specifically to segregate for novel heading date and plant height QTL. Our expanding knowledge of variants underlying these oligogenic traits results in the development of breeding populations where the major QTL will be known and predictions for heading date and plant height can be made. If we have genotypes for the causal polymorphisms underlying these QTL, we can make predictions in new populations regardless of their relationship to the training population lines. Fine-mapping and marker development for these and further novel QTL will then improve prediction models.

### Plant growth QTL and variation for source traits

In the past few years, a number of variants impacting yield component traits that generate variation in sink tissues have been identified and cloned (Wang et al., 2018; Dixon et al., 2018; Kuzay et al., 2019; DeWitt et al., 2020). However, increasing the frequency of variants associated with larger grain size and number will only increase yields if plants produce sufficient carbohydrates (“source”) to fill those grains. Similar characterization of important QTL underlying variation in physiological source traits will therefore also be critical to understand the components of yield variation. Variation in NDVI (normalized difference vegetation index) measurements or direct biomass samples, taken as proxies for source availability, is related to variation in the plant growth traits studied here. Both heading date and adult plant height can be viewed as components of the continuous phenotype of plant growth over time. While adult plant height is controlled by what are termed plant height QTL, juvenile plant height is often understood as winter dormancy release and is largely under the same genetic control as heading date (Guedira et al., 2014). To understand the genetic basis of plant growth in this population, we measured plant height over multiple days during development. We showed that variation in plant growth is influenced by a combination of heading date and plant height QTL. Studies of plant source traits may find it useful to consider phenotyping experiments for plant height and heading date as well to distinguish between QTL for heading date and plant height, and true source or biomass QTL. Understanding how plant height and heading date QTL interact to generate variation in plant growth over time will be critical to understanding how they impact source traits and in characterizing novel plant source QTL that can be deployed for higher yielding genotypes.

## Conclusions

The polygenic nature of wheat yield results in part from major and moderate QTL for adaptation traits and other phenotypes that influence yield. It is therefore useful to consider and select for component phenotypes like disease resistance and plant growth traits that can influence yield separately, and to properly model these traits we need to first understand their genetic architectures. Already the simple genetic basis of many disease resistance genes has made MAS for disease resistance in wheat very useful to breeders, a success story that could be replicated with plant growth traits given cost-effective predictions. Here, we show that component phenotypes of plant growth over time have an oligogenic basis dominated by QTL of major and moderate effect that allows for their prediction with simple QTL regression models. The movement towards genomic selection has called into question the utility of fine-mapping and positional cloning studies. We demonstrate the importance of major QTL and the poor performance of standard models in this study, illustrating the utility of understanding the important variants underlying these traits and others to crop improvement.

## Author’s contributions

ND wrote the initial draft of the manuscript and performed all analyses. ND, MG, EL, JBH, and GBG edited the manuscript. MG, EL, and GBG designed the experiment and developed the population. ND, MG, JPM, DM, MM, and JJ planted the experiment and assisted with data collection and harvest. JBH assisted with data analysis and experimental design.

## Acknowledgements

The authors thank staff of the USDA-ARS Plant Science Unit for assistance in genotyping and field evaluation of populations. Special thanks to Dr. Brian Ward, who gave valuable advice on many of the methods in this manuscript, Anna Rogers, who provided invaluable input on multiple aspects of the paper, and Kim Howell and Jared Smith, who extracted DNA and prepared GBS libraries. Support was provided by the Agriculture and Food Research Initiative Competitive Grant 67007-25939 (WheatCAP-IWYP) from the USDA NIFA.

## Additional Files

### Additional file 1 — Additional file 1.docx

Word document containing QTL results for spike length in figure S1, and KASP marker nucleotide sequences in table S1.

## Additional file 2 — Additional file 2.xlsx

Excel file containing four tables representing QTL model output and additive QTL model output for plant height and heading date in all environments.

## References

Christopher K. Addison, R. Esten Mason, Gina Brown-Guedira, Mohammed Guedira, Yuanfeng Hao, Randall G. Miller, Nithya Subramanian, Dennis N. Lozada, Andrea Acuna, Maria N. Arguello, Jerry W. Johnson, Amir M.H. Ibrahim, Russell Sutton, and Stephen A. Harrison. QTL and major genes influencing grain yield potential in soft red winter wheat adapted to the southern United States. Euphytica, 209(3):665–677, 6 2016. ISSN 15735060. doi: 10.1007/s10681-016-1650-1.

Eduard D Akhunov, Alina R Akhunova, Olin D Anderson, James A Anderson, Nancy Blake, Michael T Clegg, Devin Coleman-Derr, Emily J Conley, Curt C Crossman, Karin R Deal, Jorge Dubcovsky, Bikram S Gill, Yong Q Gu, Jakub Hadam, Hwayoung Heo, Naxin Huo, Gerard R Lazo, Ming-Cheng Luo, Yaqin Q Ma, David E Matthews, Patrick E McGuire, Peter L Morrell, Calvin O Qualset, James Renfro, Dindo Tabanao, Luther E Talbert, Chao Tian, Donna M Toleno, Marilyn L Warburton, Frank M You, Wenjun Zhang, and Jan Dvorak. Nucleotide diversity maps reveal variation in diversity among wheat genomes and chromosomes. BMC Genomics, 11(1):702, 2010. ISSN 1471-2164. doi: 10.1186/1471-2164-11-702. URL http://bmcgenomics.biomedcentral.com/articles/10.1186/1471-2164-11-702.

James Beales, Adrian Turner, Simon Griffiths, John W. Snape, and David A. Laurie. A Pseudo-Response Regulator is misexpressed in the photoperiod insensitive Ppd-D1a mutant of wheat (Triticum aestivum L.). Theoretical and Applied Genetics, 115(5):721–733, 9 2007. ISSN 00405752. doi: 10.1007/s00122-007-0603-4.

Rex Bernardo. Genomewide Selection when Major Genes Are Known. Crop Science, 54(1):68–75, 1 2014. ISSN 0011183X. doi: 10.2135/cropsci2013.05.0315. URL http://doi.wiley.com/10.2135/cropsci2013.05.0315.

A. Borner, A. J. Worland, J. Plaschke, Erika Schumann, and C. N. Law. Pleiotropic Effects of Genes for Reduced Height (Rht) and Day-Length Insensitivity (Ppd) on Yield and its Components for Wheat Grown in Middle Europe. Plant Breeding, 111(3):204–216, 10 1993. ISSN 0179-9541. doi: 10.1111/j.1439-0523.1993.tb00631.x. URL http://doi.wiley.com/10.1111/j.1439-0523.1993.tb00631.x.

K. W. Broman, H. Wu, S. Sen, and G. A. Churchill. R/qtl: QTL mapping in experimental crosses. Bioinformatics, 19(7):889–890, 5 2003. ISSN 1367-4803. doi: 10.1093/bioinformatics/btg112. URL https://academic.oup.com/bioinformatics/article-lookup/doi/10.1093/bioinformatics/btg112.

Edward S. Buckler, James B. Holland, Peter J. Bradbury, Charlotte B. Acharya, Patrick J. Brown, Chris Browne, Elhan Ersoz, Sherry Flint-Garcia, Arturo Garcia, Jeffrey C. Glaubitz, Major M. Goodman, Carlos Harjes, Kate Guill, Dallas E. Kroon, Sara Larsson, Nicholas K. Lepak, Huihui Li, Sharon E. Mitchell, Gael Pressoir, Jason A. Peiffer, Marco Oropeza Rosas, Torbert R. Rocheford, M. Cinta Romay, Susan Romero, Stella Salvo, Hector Sanchez Villeda, H. Sofia Da Silva, Qi Sun, Feng Tian, Narasimham Upadyayula, Doreen Ware, Heather Yates, Jianming Yu, Zhiwu Zhang, Stephen Kresovich, and Michael D. McMullen. The genetic architecture of maize flowering time. Science, 325(5941):714–718, 8 2009. ISSN 00368075. doi: 10.1126/science.1174276.

DG Butler, BR Cullis, AR Gilmour, BJ Gogel, and R Thompson. ASReml-R Reference Manual Version 4, 2017.

Yihua Chen, Brett F Carver, Shuwen Wang, Shuanghe Cao, and Liuling Yan. Genetic regulation of developmental phases in winter wheat. Molecular Breeding, 26:573–582, 2010. doi: 10.1007/s11032-010-9392-6.

Zhaoyan Chen, Xuejiao Cheng, Lingling Chai, Zihao Wang, Dejie Du, Zhihui Wang, Ruolin Bian, Aiju Zhao, Mingming Xin, Weilong Guo, Zhaorong Hu, Huiru Peng, Yingyin Yao, Qixin Sun, and Zhongfu Ni. Pleiotropic QTL influencing spikelet number and heading date in common wheat (Triticum aestivum L.). Theoretical Applied Genetics, page 3, 2020. doi: 10.1007/s00122-020-03556-6. URL https://doi.org/10.1007/s00122-020-03556-6.

Noah DeWitt, Mohammed Guedira, Edwin Lauer, Martin Sarinelli, Priyanka Tyagi, Daolin Fu, QunQun Hao, J. Paul Murphy, David Marshall, Alina Akhunova, Katherine Jordan, Eduard Akhunov, and Gina Brown-Guedira. Sequence-based mapping identifies a candi-date transcription repressor underlying awn suppression at the B1 locus in wheat. New Phytologist, 225(1):326–339, 1 2020. ISSN 0028-646X. doi: 10.1111/nph.16152. URL https://onlinelibrary.wiley.com/doi/abs/10.1111/nph.16152.

Aurora Díaz, Meluleki Zikhali, Adrian S. Turner, Peter Isaac, and David A. Laurie. Copy num-ber variation affecting the photoperiod-B1 and vernalization-A1 genes is associated with altered flowering time in wheat (Triticum aestivum). PLoS ONE, 7(3), 3 2012. ISSN 19326203. doi: 10.1371/journal.pone.0033234.

Laura E. Dixon, Julian R. Greenwood, Stefano Bencivenga, Peng Zhang, James Cockram, Gregory Mellers, Kerrie Ramm, Colin Cavanagh, Steve M. Swain, and Scott A. Boden. TEOSINTE BRANCHED1 regulates inflorescence architecture and development in bread wheat (Triticum aestivum). Plant Cell, 30(3):563–581, 3 2018. ISSN 1532298X. doi: 10.1105/tpc.17.00961.

RJ Elshire, JC Glaubitz, Q Sun, JA Poland, and K Kawamoto. A Robust, Simple Genotyping-by-Sequencing (GBS) Approach for High Diversity Species. PLoS ONE, 6(5):19379, 2011. doi: 10.1371/journal.pone.0019379. URL http://www.illumina.com/Documents/.

Jeffrey B. Endelman. Ridge Regression and Other Kernels for Genomic Selection with R Package rrBLUP. The Plant Genome, 4(3):250–255, 11 2011. ISSN 19403372. doi: 10.3835/plantgenome2011.08.0024. URL http://doi.wiley.com/10.3835/plantgenome2011.08.0024.

FAO. Crop Prospects and Food Situation #1, March 2020. FAO, 2020. ISBN 978-92-5-132262-8. doi: 10.4060/ca8032en. URL http://www.fao.org/documents/card/en/c/ca8032en.

R. A. Fischer and K. J. Quail. The effect of major dwarfing genes on yield potential in spring wheats. Euphytica, 46(1):51–56, 3 1990. ISSN 00142336. doi: 10.1007/BF00057618.

Daolin Fu, Péter Szűcs, Liuling Yan, Marcelo Helguera, Jeffrey S. Skinner, Jarislav Von Zitze-witz, Patrick M. Hayes, and Jorge Dubcovsky. Large deletions within the first intron in VRN-1 are associated with spring growth habit in barley and wheat. Molecular Genetics and Genomics, 273(1):54–65, 3 2005. ISSN 16174615. doi: 10.1007/s00438-004-1095-4. URL http://www.ncbi.nlm.nih.gov/pubmed/15690172.

Debora Gasperini, Andy Greenland, Peter Hedden, Rene Dreos, Wendy A Harwood, and Simon Griffiths. Genetic and physiologcial analysis of Rht8 in bread wheat: an alternative source of semi-dwarfism with a reduced sensitivty to brassinosteroids. Journal of Experimental Botany, pages 4419–4436, 2012.

Jeffrey C. Glaubitz, Terry M. Casstevens, Fei Lu, James Harriman, Robert J. Elshire, Qi Sun, and Edward S. Buckler. TASSEL-GBS: A High Capacity Genotyping by Sequencing Analysis Pipeline. PLoS ONE, 9(2):e90346, 2 2014. ISSN 1932-6203. doi: 10.1371/journal.pone.0090346. URL https://dx.plos.org/10.1371/journal.pone.0090346.

Mohammed Guedira, Peter Maloney, Mai Xiong, Stine Petersen, J. Paul Murphy, David Marshall, Jerry Johnson, Steve Harrison, and Gina Brown-Guedira. Vernalization duration requirement in soft winter wheat is associated with variation at the VRN-B1 locus. Crop Science, 54(5):1960–1971, 2014. ISSN 14350653. doi: 10.2135/cropsci2013.12.0833.

Mohammed Guedira, Mai Xiong, Yuan Feng Hao, Jerry Johnson, Steve Harrison, David Marshall, and Gina Brown-Guedira. Heading date QTL in winter wheat (Triticum aestivum L.) coincide with major developmental genes VERNALIZATION1 and PHOTOPERIOD1. PLoS ONE, 11(5), 5 2016. ISSN 19326203. doi: 10.1371/journal.pone.0154242.

Peter Hedden. The genes of the Green Revolution, 1 2003. ISSN 01689525.

Ning Huang, Brigitte Courtois, Gurdev S. Khush, Hongxuan Lin, Guoliang Wang, Ping Wu, and Kangle Zheng. Association of quantitative trait loci for plant height with major dwarfing genes in rice. Heredity, 77(2):130–137, 1996. ISSN 0018067X. doi: 10.1038/hdy.1996.117.

Nestor Kippes, Mohammed Guedira, Lijuan Lin, Maria A. Alvarez, Gina L. Brown-Guedira, and Jorge Dubcovsky. Single nucleotide polymorphisms in a regulatory site of VRN-A1 first intron are associated with differences in vernalization requirement in winter wheat. Molecular Genetics and Genomics, 293(5):1231–1243, 10 2018. ISSN 16174623. doi: 10.1007/s00438-018-1455-0.

V. Korzun, M. S. Röder, M. W. Ganal, A. J. Worland, and C. N. Law. Genetic analysis of the dwarfing gene (Rht8) in wheat. Part I. Molecular mapping of Rht8 on the short arm of chromosome 2D of bread wheat (Triticum aestivum L.). Theoretical and Applied Genetics, 96(8):1104–1109, 6 1998. ISSN 00405752. doi: 10.1007/s001220050845.

Saarah Kuzay, Yunfeng Xu, Junli Zhang, Andrew Katz, Stephen Pearce, Zhenqi Su, Max Fraser, James A. Anderson, Gina Brown-Guedira, Noah DeWitt, Amanda Peters Haugrud, Justin D. Faris, Eduard Akhunov, Guihua Bai, and Jorge Dubcovsky. Identification of a candidate gene for a QTL for spikelet number per spike on wheat chromosome arm 7AL by high-resolution genetic mapping. Theoretical and Applied Genetics, 132(9):2689–2705, 9 2019. ISSN 14322242. doi: 10.1007/s00122-019-03382-5.

Genqiao Li, Ming Yu, Tilin Fang, Shuanghe Cao, Brett F. Carver, and Liuling Yan. Vernalization requirement duration in winter wheat is controlled by TaVRN-A1 at the protein level. Plant Journal, 76(5):742–753, 12 2013. ISSN 09607412. doi: 10.1111/tpj.12326.

Heng Li, Bob Handsaker, Alec Wysoker, Tim Fennell, Jue Ruan, Nils Homer, Gabor Marth, Goncalo Abecasis, and Richard Durbin. The Sequence Alignment/Map format and SAMtools. Bioinformatics, 25(16):2078–2079, 8 2009. ISSN 13674803. doi: 10.1093/bioinformatics/btp352.

Shinichi Nakagawa and Holger Schielzeth. A general and simple method for obtaining R2 from generalized linear mixed-effects models. Methods in Ecology and Evolution, 4 (2):133–142, 2 2013. ISSN 2041210X. doi: 10.1111/j.2041-210x.2012.00261.x. URL http://doi.wiley.com/10.1111/j.2041-210x.2012.00261.x.

Hidetaka Nishida, Tetsuya Yoshida, Kohei Kawakami, Masaya Fujita, Bo Long, Yukari Akashi, David A. Laurie, and Kenji Kato. Structural variation in the 5 upstream region of photoperiod-insensitive alleles Ppd-A1a and Ppd-B1a identified in hexaploid wheat (Triticum aestivum L.), and their effect on heading time. Molecular Breeding, 31(1):27–37, 7 2013. ISSN 13803743. doi: 10.1007/s11032-012-9765-0.

Kazusa Nishimura, Ryuji Moriyama, Keisuke Katsura, Hiroki Saito, Rihito Takisawa, Akira Kitajima, and Tetsuya Nakazaki. The early flowering trait of an emmer wheat accession (Triticum turgidum L. ssp. dicoccum) is associated with the cis-element of the Vrn-A3 locus. Theoretical and Applied Genetics, 131(10):2037–2053, 10 2018. ISSN 00405752. doi: 10.1007/s00122-018-3131-5.

Jason A. Peiffer, Maria C. Romay, Michael A. Gore, Sherry A. Flint-Garcia, Zhiwu Zhang, Mark J. Millard, Candice A.C. Gardner, Michael D. McMullen, James B. Holland, Peter J. Bradbury, and Edward S. Buckler. The genetic architecture of maize height. Genetics, 196(4):1337–1356, 4 2014. ISSN 19432631. doi: 10.1534/genetics.113.159152.

Paulino Pérez and Gustavo De Los Campos. Genome-Wide Regression and Prediction with the BGLR Statistical Package. Genetics, 198:483, 2014. doi: 10.1534/genetics.114.164442. URL http://www.genetics.org/lookup/suppl/.

Jesse Poland, Jeffrey Endelman, Julie Dawson, Jessica Rutkoski, Shuangye Wu, Yann Manes, Susanne Dreisigacker, José Crossa, Héctor Sánchez-Villeda, Mark Sorrells, and Jean Luc Jannink. Genomic selection in wheat breeding using genotyping-by-sequencing. Plant Genome, 5(3):103– 113, 2012. ISSN 19403372. doi: 10.3835/plantgenome2012.06.0006.

G. J. Rebetzke and R. A. Richards. Gibberellic acid-sensitive dwarfing genes reduce plant height to increase kernel number and grain yield of wheat. Australian Journal of Agricultural Research, 51 (2):235–245, 2000. ISSN 00049409. doi: 10.1071/AR99043.

G. J. Rebetzke, M. H. Ellis, D. G. Bonnett, B. Mickelson, A. G. Condon, and R. A. Richards. Height reduction and agronomic performance for selected gibberellin-responsive dwarfing genes in bread wheat (Triticum aestivum L.). Field Crops Research, 126:87–96, 2 2012. ISSN 03784290. doi: 10.1016/j.fcr.2011.09.022.

Trevor W. Rife and Jesse A. Poland. Field book: An open-source application for field data collection on android, 2014. ISSN 14350653.

J. Martin Sarinelli, J. Paul Murphy, Priyanka Tyagi, James B. Holland, Jerry W. Johnson, Mohamed Mergoum, Richard E. Mason, Ali Babar, Stephen Harrison, Russell Sutton, Carl A. Griffey, and Gina Brown-Guedira. Training population selection and use of fixed effects to optimize genomic predictions in a historical USA winter wheat panel. Theoretical and Applied Genetics, 132(4):1247–1261, 4 2019. ISSN 00405752. doi: 10.1007/s00122-019-03276-6.

Lindsay M Shaw, Bo Lyu, Rebecca Turner, Chengxia Li, Fengjuan Chen, Xiuli Han, Daolin Fu, and Jorge Dubcovsky. FLOWERING LOCUS T2 regulates spike development and fertility in temperate cereals. Journal of Experimental Botany, 70(1):193–204, 1 2019. ISSN 0022-0957. doi: 10.1093/jxb/ery350. URL https://academic.oup.com/jxb/article/70/1/193/5118414.

Julian Taylor and David Butler. R package ASMap: Efficient genetic linkage map construction and diagnosis. Journal of Statistical Software, 79, 2017. ISSN 15487660. doi: 10.18637/jss.v079.i06.

Wei Wang, James Simmonds, Qianli Pan, Dwight Davidson, Fei He, Abdulhamit Battal, Alina Akhunova, Harold N. Trick, Cristobal Uauy, and Eduard Akhunov. Gene editing and mutagenesis reveal inter-cultivar differences and additivity in the contribution of TaGW2 homoeologues to grain size and weight in wheat. Theoretical and Applied Genetics, 131(11):2463–2475, 11 2018. ISSN 00405752. doi: 10.1007/s00122-018-3166-7.

Brian P. Ward, Gina Brown-Guedira, Priyanka Tyagi, Frederic L. Kolb, David A. Van Sanford, Clay H. Sneller, and Carl A. Griffey. Multienvironment and Multitrait Genomic Selection Models in Unbalanced Early-Generation Wheat Yield Trials. Crop Science, 59(2):491–507, 3 2019. ISSN 0011183X. doi: 10.2135/cropsci2018.03.0189. URL http://doi.wiley.com/10.2135/cropsci2018.03.0189.

A. J. Worland, V. Korzun, M. S. Röder, M. W. Ganal, and C.N. Law. Genetic analysis of the dwarfing gene Rht8 in wheat. Part II. The distribution and adaptive significance of allelic variants at the Rht8 locus of wheat as revealed by microsatellite screening. Theoretical and Applied Genetics, 96(8):1110–1120, 6 1998. ISSN 00405752. doi: 10.1007/s001220050846.

L. Yan, M. Helguera, K. Kato, S. Fukuyama, J. Sherman, and J. Dubcovsky. Allelic variation at the VRN-1 promoter region in polyploid wheat. Theoretical and Applied Genetics, 109(8):1677–1686, 11 2004. ISSN 00405752. doi: 10.1007/s00122-004-1796-4.

L. Yan, D. Fu, C. Li, A. Blechl, G. Tranquilli, M. Bonafede, A. Sanchez, M. Valarik, S. Yasuda, and J. Dubcovsky. The wheat and barley vernalization gene VRN3 is an orthologue of FT. Proceedings of the National Academy of Sciences of the United States of America, 103(51):19581–19586, 12 2006. ISSN 00278424. doi: 10.1073/pnas.0607142103.

S. Youssefian, E. J.M. Kirby, and M. D. Gale. Pleiotropic effects of the GA-insensitive Rht dwarfing genes in wheat. 2. Effects on leaf, stem, ear and floret growth. Field Crops Research, 28(3):191–210, 1 1992. ISSN 03784290. doi: 10.1016/0378-4290(92)90040-G.

Andrey Ziyatdinov, Miquel Vázquez-Santiago, Helena Brunel, Angel Martinez-Perez, Hugues Aschard, and Jose Manuel Soria. lme4qtl: linear mixed models with flexible covariance structure for genetic studies of related individuals. BMC Bioinformatics, 2018. doi: 10.1186/s12859-018-2057-x. URL https://doi.org/10.1186/s12859-018-2057-x.

