## Additional file 1 for "Genetic variation for plant growth traits in a common wheat population is dominated by known variants and novel QTL"

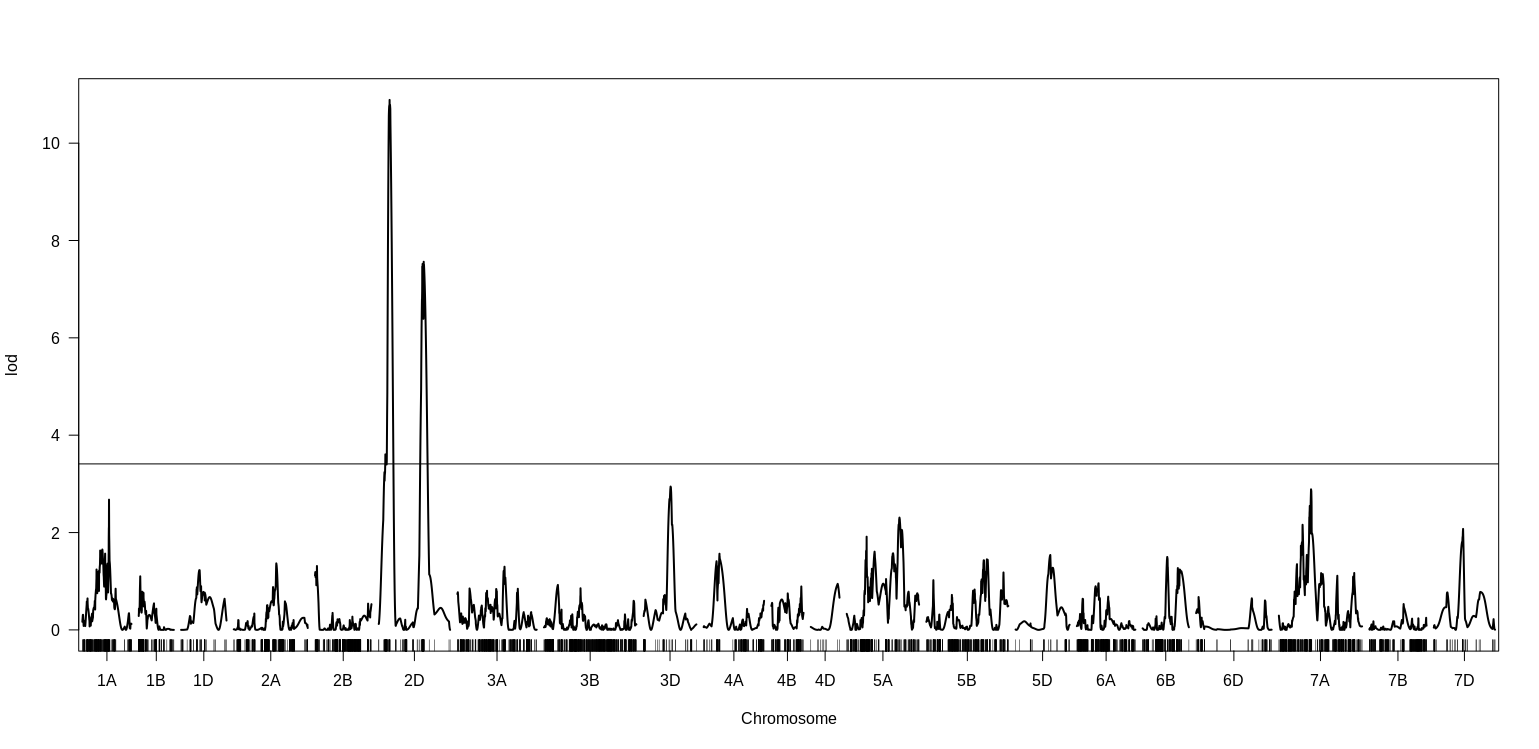

**Figure S1.** CIM QTL results for spike compactness in Raleigh 2019. Spike compactness was rated visually on a 1-5 scale, with 5 being most compact with a club wheat-like phenotype. Significance declared at a LOD of 3.41 for alpha = .05 based on a thousand permutations. The heading date locus on the short arm of chromosome 2D colocates with the major QTL, suggesting that it is likely *Rht8*.

| Marker_name | Chr | Sequence |
| --- | --- | --- |
| Rht-D1_A1 | 4D | GAAGGTGACCAAGTTCATGCTCATGGCCATCTCGAGCTRCTC |
| Rht-B1_A2 | 4D | GAAGGTCGGAGTCAACGGATTCATGGCCATCTCGAGCTRCTA |
| Rht-D1_C | 4D | CGGGTACAAGGTGCGCGCC |
| Ppd-D1_A1 | 2D | GAAGGTCGGAGTCAACGGATTAAGAGGAAACATGTTGGGGTCC |
| Ppd-D1_A2 | 2D | GAAGGTGACCAAGTTCATGCTCAAGGAAGTATGAGCAGCGGTT |
| Ppd-D1_C | 2D | GCCTCCCACTACACTGGGC |
| FTA2_A1 | 3A | GAAGGTGACCAAGTTCATGCTACGTCCACCGGCATCTTGGAC |
| FTA2_A2 | 3A | GAAGGTCGGAGTCAACGGATTCGTCCACCGGCATCTTGGAT |
| FTA2_C | 3A | GTACAGCTTCGGGGTACTGCTGTT |
| VrnA3_proDel_A1 | 7A | GAAGGTGACCAAGTTCATGCTCAGCTTACGCTTACTCTTGCTCCC |
| VrnA3_proDel_A2 | 7A | GAAGGTCGGAGTCAACGGATTCAGCTTACGCTTACTCTTGCTCCA |
| VrnA3_proDel_C | 7A | CTCCCGGCCATTTCCCCTTCC |
| B1_A1 | 5A | GAAGGTGACCAAGTTCATGCTAGCTACGGGCCCACTTRGACA |
| B1_A2 | 5A | GAAGGTCGGAGTCAACGGATTCTACGGGCCCACTTRGACG |
| B1_C1 | 5A | CCTGCGGGGCTCCCAGCAA |
| WAPOA1_Hap13vs2_A1 | 7A | GAAGGTGACCAAGTTCATGCTCTGATTATGGGCGGTTGATCTGC |
| WAPOA1_Hap13vs2_A2 | 7A | GAAGGTCGGAGTCAACGGATTTCTGATTATGGGCGGTTGATCTGT |
| WAPOA1_Hap13vs2_C1 | 7A | GACCAGCGCCGGCGACT |

**Table S1.** KASP marker sequences for variants that are tightly linked to or underly mapped QTL in this study.
